## Supplementary Tables for "*de novo* diploid genome assembly using long noisy reads"

**Supplementary Table 1. Performance of supporting read selection method of PECAT**

| Sample | *S. cerevisiae* SK1 × Y12 (CLR, 200X) | *A. thaliana* Col-0 × Cvi-0 (CLR, 164X) | *D. melanogaster* ISO1 × A4 (CLR, 200X) | *B. taurus* Angus × Brahman (CLR, 135X) | *A. thaliana* Col-0 × C24 (ONT, 106X) | *B. taurus* Bison × Simmental (ONT, 200X) | HG002 (ONT, R9, 59X) |
|---|---|---|---|---|---|---|---|
| Heterozygosity rate (%) | 0.85 | 1.04 | 0.84 | 1.12 | 0.83 | 1.48 | 0.34 |
| Percentage of selected reads in all supporting reads (%) | 56.8 | 63.9 | 54.8 | 54.5 | 56.7 | 53.6 | 56.5 |
| Percentage of inconsistent reads in all supporting reads (%) | 36.7 | 31.5 | 41.1 | 28.1 | 33.8 | 37.2 | 27.4 |
| Percentage of inconsistent reads in selected reads (%) | 2.8 | 3.5 | 4.9 | 3.6 | 4.3 | 2.9 | 4.2 |
| Percentage of inconsistent reads in selected reads after re-weighting the scores (%) | 2.1 | 3.1 | 4.0 | 3.1 | 3.5 | 2.3 | 3.4 |

'CLR' indicates there are PacBio CLR reads. 'ONT' indicates there are Nanopore reads. 'R9' indicates it is Nanopore R9 data.

**Supplementary Table 2. Performance comparison of error correction on simulated data**

| Sample | Pipeline | Size (Mb) | Max (Kb) | N50 (Kb) | Accuracy of base (%) | Accuracy of SNP (%) |
|---|---|---|---|---|---|---|
| PacBio CLR het = 0.01 | raw data | 89.2 | 62.1 | 26.3 | 88.58 | 96.55 |
| | Canu | 81.5 | 56.4 | 25.6 | 99.44 | 80.53 |
| | MECAT2 | 86.3 | 60.1 | 25.8 | 99.28 | 66.84 |
| | PECAT | 87.3 | 61.0 | 25.7 | 99.77 | 99.83 |
| PacBio CLR het = 0.005 | raw data | 89.2 | 62.9 | 26.5 | 88.52 | 96.48 |
| | Canu | 81.2 | 56.6 | 25.7 | 99.54 | 76.70 |
| | MECAT2 | 86.1 | 62.1 | 26.0 | 99.45 | 59.36 |
| | PECAT | 87.3 | 61.9 | 25.9 | 99.77 | 99.84 |
| PacBio CLR het = 0.001 | raw data | 89.2 | 58.3 | 26.4 | 88.66 | 96.44 |
| | Canu | 82.2 | 54.5 | 25.8 | 99.66 | 73.29 |
| | MECAT2 | 86.5 | 57.5 | 25.9 | 99.63 | 54.27 |
| | PECAT | 87.3 | 57.3 | 25.8 | 99.76 | 99.81 |
| PacBio CLR het = 0.0005 | raw data | 89.2 | 59.0 | 26.3 | 88.64 | 96.57 |
| | Canu | 82.6 | 55.2 | 25.6 | 99.66 | 72.83 |
| | MECAT2 | 86.6 | 58.1 | 25.8 | 99.66 | 54.05 |
| | PECAT | 87.3 | 57.9 | 25.7 | 99.75 | 99.46 |
| PacBio CLR het = 0.0001 | raw data | 89.2 | 57.8 | 26.6 | 88.61 | 96.82 |
| | Canu | 82.2 | 52.2 | 25.9 | 99.67 | 72.31 |
| | NECAT | 86.4 | 56.7 | 26.1 | 99.67 | 54.64 |
| | PECAT | 87.3 | 56.5 | 26.0 | 99.78 | 91.83 |
| Nanopore het = 0.01 | raw data | 89.2 | 57.6 | 26.1 | 88.59 | 98.14 |
| | Canu | 79.2 | 53.6 | 25.4 | 99.28 | 76.41 |
| | NECAT | 87.1 | 56.8 | 25.6 | 99.40 | 58.65 |
| | PECAT | 87.3 | 56.7 | 25.5 | 99.44 | 99.94 |
| Nanopore het = 0.005 | raw data | 89.2 | 55.2 | 26.1 | 89.10 | 98.09 |
| | Canu | 80.1 | 50.7 | 25.6 | 99.69 | 73.75 |
| | NECAT | 87.2 | 54.2 | 25.7 | 99.77 | 56.67 |
| | PECAT | 87.3 | 54.2 | 25.6 | 99.94 | 99.95 |
| Nanopore het = 0.001 | raw data | 89.2 | 59.9 | 26.5 | 89.06 | 98.22 |
| | Canu | 79.9 | 53.1 | 25.9 | 99.79 | 72.73 |
| | NECAT | 87.2 | 57.9 | 26.0 | 99.87 | 56.86 |
| | PECAT | 87.3 | 57.8 | 25.9 | 99.93 | 99.90 |
| Nanopore het = 0.0005 | raw data | 89.2 | 58.1 | 26.6 | 89.10 | 98.07 |
| | Canu | 80.2 | 52.0 | 26.0 | 99.69 | 71.64 |
| | NECAT | 87.3 | 57.2 | 26.1 | 99.77 | 55.96 |
| | PECAT | 87.3 | 57.0 | 26.0 | 99.94 | 99.72 |
| Nanopore het = 0.0001 | raw data | 89.2 | 62.1 | 26.3 | 89.05 | 98.25 |
| | Canu | 79.5 | 52.6 | 25.7 | 99.82 | 73.09 |

| | | | | | | |
|---|---|---|---|---|---|---|
| | NECAT | 87.4 | 61.3 | 25.8 | 99.87 | 54.81 |
| | PECAT | 87.3 | 61.3 | 25.7 | 99.95 | 94.10 |

The simulated data is illustrated in **Supplementary Note 1**. ‘Size’ is the total number of base pairs of the reads. ‘Max’ is the max length of the reads. ‘N50’ is the length of the shortest read for which longer and equal-length reads cover at least 50% of the corrected reads. ‘Accuracy of base’ is the percentage of matched base pairs in the alignments between the reads and the reference genome. ‘Accuracy of SNP’ is the accuracy of SNP alleles in reads. ‘het’ represents the heterozygosity rate of the simulated data.

**Supplementary Table 3. Intermediate state of the first round of assembly**

| Sample | Symbol / Formula | *S. cerevisiae* SK1 × Y12 | *A. thaliana* Col-0 × Cvi-0 | *D. melanogaster* ISO1 × A4 | *B. taurus* Angus × Brahman | *A. thaliana* Col-0 × C24 | *B. taurus* Bison × Simmental | HG002 |
|---|---|---|---|---|---|---|---|---|
| Number of all candidate overlaps | N1 | 15765862 | 217154507 | 126571948 | 451388849 | 761657318 | 470660333 | 529956421 |
| Number of overlaps without long overhang | N2 | 14931099 | 194873544 | 22549502 | 393535386 | 609033764 | 190990689 | 231772901 |
| Number of the overlaps for constructing the string graph after removing the overlaps whose reads are contained in other reads or with low coverage | N3 | 21689 | 392687 | 193075 | 3376177 | 197563 | 2350982 | 344684 |
| Number of all edges in the string graph* | 2*N3 | 43378 | 785374 | 386150 | 6752354 | 395126 | 4701964 | 689368 |
| Number of the transitive edges in the string graph | N4 | 32486 | 654336 | 323358 | 5502574 | 313326 | 3978100 | 526502 |
| Number of low-quality edges in the string graph | N5 | 234 | 15980 | 5942 | 28094 | 10814 | 13340 | 11182 |
| Number of transitive edges which are reactivated in the string graph | N6 | 218 | 802 | 476 | 5094 | 412 | 2164 | 850 |
| Number of the overlaps introduced by contained reads | N7 | 64 | 1527 | 337 | 3830 | 1525 | 3063 | 1548 |
| Percentage of the overlaps with long overhang in all candidate overlaps | 1-N2/N1 | 5.3% | 10.3% | 82.2% | 12.8% | 20.0% | 59.4% | 56.3% |
| Percentage of the overlaps for constructing the string graph in the overlaps without long overhang | N3/N2 | 0.15% | 0.18% | 0.86% | 0.86% | 0.03% | 1.23% | 0.15% |
| Percentage of active edges in all edges in the string graph | 1-N4/(2*N3) | 25.1% | 16.7% | 16.3% | 18.5% | 20.7% | 15.4% | 23.6% |
| Percentage of low-quality edges in active edges in the string graph | N5/(2*N3-N4) | 2.1% | 12.2% | 9.5% | 2.2% | 13.2% | 1.8% | 6.9% |
| Percentage of transitive edges that are reactivated in the string graph | N6/N4 | 0.67% | 0.12% | 0.15% | 0.09% | 0.13% | 0.05% | 0.16% |

| | | | | | | | | |
|---|---|---|---|---|---|---|---|---|
| Percentage of edges introduced by contained reads | N7/N3 | 0.30% | 0.39% | 0.17% | 0.11% | 0.77% | 0.13% | 0.45% |

*An overlap corresponds to two edges in the string graph. ‘Symbol / Formula’ is a symbol for the corresponding metric or a formula to calculate the corresponding metric.

**Supplementary Table 4. Performance of inconsistent overlaps identification method of PECAT**

| Sample | Reads used | ASM1 (%) | ASM2 (%) |
|---|---|---|---|
| *S. cerevisiae*, SK1 × Y12, CLR | Corrected | 36.98 | 0.32 |
| *A. thaliana*, Col-0 × Cvi-0, CLR | Corrected | 16.00 | 0.57 |
| *D. melanogaster*, ISO1 × A4, CLR | Corrected | 15.94 | 0.90 |
| *B. taurus*, Angus × Brahman, CLR | Corrected | 36.55 | 0.63 |
| *A. thaliana*, Col-0 × C24, ONT | Corrected | 16.62 | 1.57 |
| | Raw | 16.62 | 1.57 |
| | Corrected +Raw | 16.62 | 0.79 |
| *B. taurus*, Bison × Simmental, ONT | Corrected | 18.08 | 0.06 |
| | Raw | 18.08 | 0.07 |
| | Corrected +Raw | 18.08 | 0.03 |
| HG002, ONT, R9 | Corrected | 37.45 | 1.46 |
| | Raw | 37.45 | 1.12 |
| | Corrected+Raw | 37.45 | 1.09 |

'ASM1' and 'ASM2' are the percentages of inconsistent overlaps in simplified assembly graphs in the first and the second round of assembly. 'CLR' indicates there are PacBio CLR reads. 'ONT' indicates there are Nanopore reads. 'R9' indicates it is Nanopore R9 data. 'Reads used' means which reads are used to identify inconsistent overlaps. 'Corrected' and 'Raw' represent corrected reads and raw reads, respectively.

**Supplementary Table 5. Performance of assemblies before and after removing inconsistent overlaps of PECAT.**

| | The first round of assembly | | | | The second round of assembly | | | |
|---|---|---|---|---|---|---|---|---|
| Sample | Size (Mb) | Count | Max (Mb) | NG50 (Mb) | Size (Mb) | Count | Max (Mb) | NG50 (Mb) |
| *S. cerevisiae* | 12.9 | 83 | 1.48 | 0.89 | 12.3 | 24 | 1.5 | 0.8 |
| SK1×Y12, CLR | 0.9 | 27 | 0.06 | 0.00 | 11.8 | 33 | 1.5 | 0.8 |
| *A. thaliana* | 139.8 | 489 | 16.84 | 14.07 | 130.6 | 373 | 16.6 | 14.3 |
| Col-0×Cvi-0, CLR | 74.0 | 845 | 0.58 | 0.05 | 120.4 | 163 | 12.8 | 7.8 |
| *D. melanogaster* | 164.2 | 391 | 28.1 | 23.4 | 149.6 | 243 | 27.1 | 24.5 |
| ISO1×A4, CLR | 84.5 | 408 | 1.8 | 0.1 | 135.7 | 201 | 27.0 | 11.9 |
| *B. taurus* | 2762.6 | 3397 | 110.8 | 55.2 | 2744.7 | 2445 | 157.2 | 72.4 |
| Angus×Brahman, CLR | 128.8 | 1940 | 0.5 | 0.0 | 2447.6 | 3424 | 17.6 | 2.8 |
| *A. thaliana* | 138.6 | 308 | 15.3 | 13.6 | 131.1 | 136 | 16.8 | 14.3 |
| Col-0×C24, ONT | 63.7 | 439 | 1.7 | 0.0 | 123.8 | 95 | 14.8 | 7.7 |
| *B. taurus* | 2984.2 | 1092 | 156.3 | 72.9 | 2963.2 | 498 | 161.1 | 94.1 |
| Bison×Simmental, ONT | 1519.8 | 5587 | 1.8 | 0.2 | 2765.5 | 1113 | 158.9 | 93.6 |
| HG002 | 3125.3 | 735 | 169.9 | 85.6 | 3058.7 | 200 | 153.3 | 92.9 |
| ONT, R9 | 137.5 | 267 | 2.6 | 0.0 | 2857.2 | 418 | 49.4 | 15.0 |

‘Size’ is the total number of base pairs in all contigs generated by PECAT. ‘Max’ is the max length of the contigs. ‘NG50’ is the length of the shortest contig for which longer and equal length contigs cover at least 50% of genome size. The genome sizes of *S. cerevisiae*, *A. thaliana*, *D. melanogaster*, *B. taurus*, and HG002 that we used for evaluation are 12M, 130M, 140M, 2.7G, and 3G, respectively. The primary and alternate contigs are separately reported on the top and the bottom of each row.

**Supplementary Table 6. Performance comparison of error correction**

| Sample | Pipeline | N50 | Match (%) | Mismatch (%) | Insertion (%) | Deletion (%) | Consistency (%) | Completeness PAT (%) / MAT (%) |
|---|---|---|---|---|---|---|---|---|
| *S. cerevisiae* (SK1 × Y12, CLR, 200X) | raw data | 13988 | 87.80 | 1.91 | 7.32 | 2.97 | 93.4 | 37.7 / 36.6 |
| | Canu | 11606 | 99.53 | 0.11 | 0.16 | 0.20 | 89.8 | 94.3 / 95.9 |
| | FALCON | 10400 | 99.69 | 0.08 | 0.03 | 0.20 | 88.5 | 73.3 / 82.2 |
| | MECAT2 | 11734 | 99.37 | 0.09 | 0.46 | 0.08 | 87.4 | 90.5 / 88.0 |
| | PECAT | 11785 | 99.71 | 0.02 | 0.12 | 0.15 | 99.7 | 98.7 / 99.2 |
| *A. thaliana* (Col-0 × Cvi-0, CLR, 164X) | raw data | 26088 | 88.24 | 2.06 | 5.54 | 4.16 | 86.6 | 40.3 / 41.0 |
| | Canu | 23105 | 99.31 | 0.14 | 0.16 | 0.38 | 94.8 | 97.6 / 98.2 |
| | FALCON | 20530 | 99.39 | 0.11 | 0.10 | 0.40 | 92.7 | 89.9 / 90.5 |
| | MECAT2 | 23285 | 99.27 | 0.13 | 0.48 | 0.12 | 94.4 | 96.5 / 97.1 |
| | PECAT | 23414 | 99.44 | 0.15 | 0.13 | 0.28 | 99.7 | 98.9 / 99.4 |
| *D. melanogaster* (ISO1 × A4, CLR, 200X) | raw data | 60067 | 89.58 | 2.72 | 5.14 | 2.56 | 93.6 | 63.1 / 64.8 |
| | Canu | 54896 | 99.45 | 0.11 | 0.24 | 0.20 | 91.4 | 93.5 / 97.2 |
| | FALCON | 37210 | 98.90 | 0.05 | 0.28 | 0.76 | 93.4 | 90.7 / 93.8 |
| | MECAT2 | 47894 | 99.01 | 0.08 | 0.77 | 0.14 | 90.0 | 91.5 / 94.6 |
| | PECAT | 51149 | 99.57 | 0.03 | 0.23 | 0.17 | 99.9 | 94.7 / 98.6 |
| *B. taurus* (Angus × Brahman, CLR, 135X) | raw data | 26938 | 86.25 | 3.09 | 7.02 | 3.64 | 87.7 | 12.9 / 12.1 |
| | MECAT2 | 21517 | 98.92 | 0.12 | 0.80 | 0.17 | 85.3 | 88.5 / 88.9 |
| | PECAT | 21760 | 99.40 | 0.07 | 0.18 | 0.35 | 99.7 | 96.9 / 97.2 |
| *A. thaliana* (Col-0 × C24, ONT, 106X) | raw data | 30858 | 92.33 | 2.79 | 1.81 | 3.06 | 95.4 | 90.1 / 89.4 |
| | Canu | 30092 | 98.64 | 0.39 | 0.15 | 0.82 | 89.1 | 90.4 / 89.6 |
| | NECAT | 29551 | 98.95 | 0.32 | 0.29 | 0.45 | 92.4 | 88.8 / 88.0 |
| | PECAT | 29799 | 99.00 | 0.39 | 0.15 | 0.46 | 99.4 | 96.7 / 96.0 |
| *B. taurus* (Bison × Simmental, ONT, 200X) | raw data | 83396 | 89.12 | 3.58 | 2.68 | 4.62 | 93.7 | 69.4 / 69.4 |
| | NECAT | 82988 | 98.78 | 0.31 | 0.36 | 0.55 | 85.7 | 80.2 / 80.9 |
| | PECAT | 80587 | 98.61 | 0.26 | 0.20 | 0.92 | 99.3 | 92.2 / 93.5 |
| HG002 (ONT, R9 59X) | raw data | 107537 | 94.97 | 1.81 | 1.24 | 1.97 | 98.8 | 98.6 / 98.6 |
| | NECAT | 106400 | 99.35 | 0.35 | 0.17 | 0.12 | 91.1 | 74.8 / 64.8 |
| | PECAT | 104009 | 99.85 | 0.04 | 0.04 | 0.08 | 99.9 | 98.8 / 99.1 |

'Match', 'Mismatch', 'Insertion', and 'Deletion' are the percentages of matches, mismatches, insertions, and deletions in the alignments between the reads and the reference genome. 'Size' is the total number of base pairs of the corrected reads. 'N50' is the length of the shortest read for which longer and equal-length reads cover at least 50% of the 40X longest reads. 'Consistency' is defined as $\sum \max(k_p, k_m)/\sum(k_p + k_m)$, in which $k_p$ and $k_m$ are the number of paternal and maternal haplotype-specific k-mers in each read. 'Completeness' is the percentage of parent-specific k-mers (occurrences $\geq 4$) in the 40X longest reads. 'PAT' and 'MAT' are paternal Completeness and maternal Completeness respectively.

**Supplementary Table 7. Accuracy of raw reads and corrected reads by NECAT and PECAT in difficult-to-map regions and low-complexity regions of HG002 reference genome**

| Region | raw data | NECAT | PECAT |
|---|---|---|---|
| Segmental duplications | 94.83% | 98.90% | 99.79% |
| Low-mappability regions | 94.94% | 98.77% | 99.57% |
| 250-bp+ non-unique regions | 94.45% | 98.63% | 99.43% |
| Homopolymer (7-11 bp) | 93.77% | 98.97% | 99.44% |
| Homopolymer (>11 bp) | 93.27% | 97.61% | 98.54% |
| Dimer repeat regions (11-50bp) | 94.39% | 98.75% | 99.52% |
| Trimer repeat regions (15-50 bp) | 94.96% | 99.21% | 99.77% |

**Supplementary Table 8. Precision, recall, and F1-score of small variants (SNP, INDEL) and structural variants (SV) in HG002 assemblies.**

| Pipeline | SNP | | | INDEL | | | SV | | |
|---|---|---|---|---|---|---|---|---|---|
| | Precision (%) | Recall (%) | F1-score (%) | Precision (%) | Recall (%) | F1-score (%) | Precision (%) | Recall (%) | F1-score (%) |
| Hifiasm (pri/alt) | 99.8 | 92.4 | 96.0 | 97.9 | 90.4 | 94.0 | 92.6 | 98.1 | 95.3 |
| Hifiasm (dual) | 99.6 | 99.0 | 99.3 | 97.4 | 97.1 | 97.3 | 94.5 | 97.8 | 96.1 |
| Flye+Hapdup (dual) | 97.7 | 99.1 | 98.4 | 23.4 | 68.8 | 34.9 | 94.2 | 97.7 | 95.9 |
| Shasta (pri/alt) | 90.0 | 79.9 | 84.7 | 18.9 | 44.1 | 26.5 | 93.6 | 86.7 | 90.0 |
| PECAT (pri/alt) | 96.4 | 98.6 | 97.5 | 19.3 | 61.1 | 29.3 | 93.9 | 98.1 | 96.0 |
| PECAT (dual) | 96.4 | 98.5 | 97.4 | 19.3 | 61.1 | 29.3 | 94.2 | 98.0 | 96.0 |

The assemblies by Hifiasm are from HiFi reads. The assemblies by other methods are from Nanopore R9 reads. 'pri/alt' represents primary/alternate format. 'dual' represents dual assembly format.

**Supplementary Table 10. Running time of error correction methods.**

| Dataset | Pipeline | Size (G) | Time (H) | Speed (G/H) |
|---|---|---|---|---|
| *S. cerevisiae* SK1×Y12, CLR, 200X | Canu | 1.11 | 70 | 0.016 |
| | MECAT2 | 1.72 | 6 | 0.300 |
| | PECAT | 0.75 | 7 | 0.111 |
| *A. thaliana* Col-0 × Cvi-0, CLR, 164X | Canu | 12.41 | 712 | 0.017 |
| | MECAT2 | 17.89 | 134 | 0.133 |
| | PECAT | 8.30 | 90 | 0.093 |
| *D. melanogaster* ISO1 × A4, CLR, 200X | Canu | 13.30 | 1497 | 0.009 |
| | MECAT2 | 23.12 | 223 | 0.104 |
| | PECAT | 9.28 | 98 | 0.095 |
| *B. taurus* Angus × Brahman, CLR, 135X | MECAT2 | 232.05 | 5578 | 0.042 |
| | PECAT | 151.13 | 3254 | 0.046 |
| *A. thaliana* Col-0 × C24, ONT, 106X | Canu | 11.21 | 726 | 0.015 |
| | NECAT | 9.27 | 75 | 0.123 |
| | PECAT | 9.93 | 109 | 0.091 |
| *B. taurus* Bison × Simmental, ONT, 200X | NECAT | 217.00 | 13524 | 0.016 |
| | PECAT | 205.74 | 4217 | 0.049 |
| HG002, ONT, R9, 59X | NECAT | 171.99 | 5637 | 0.031 |
| | PECAT | 165.32 | 4404 | 0.038 |

‘Size’ is the total number of base pairs of the corrected reads. ‘Time’ is the running time of the error correction method, and the ‘Speed’ is defined as ‘Size’/’Time’. All the pipelines are tested on the same computer with a 2.0 GHz CPU and 2T GB RAM of memory and run with 48 threads. For *B. taurus* and HG002, Canu didn’t correct the raw reads in 3 weeks, so it is excluded. Since FALCON does not have an independent error correction step, the table doesn’t contain its running times.

**Supplementary Table 11. Running time, memory usage, and disk space usage of assemblers.**

| Dataset | Pipeline | CPU time (H) | Peak memory usage (G) | Peak disk space usage (G) |
|---|---|---|---|---|
| *S. cerevisiae* SK1×Y12, CLR, 200X | Canu | 109 | 7 | - |
| | FALCON-Unzip | 199 | 7 | - |
| | PECAT | 11 | 18 | 4 |
| *A. thaliana* Col-0×Cvi-0, CLR, 135X | Canu | 1456 | 12 | - |
| | FALCON-Unzip | 2652 | 35 | - |
| | PECAT | 167 | 71 | 80 |
| *D. melanogaster* ISO1×A4, CLR, 164X | Canu | 3006 | 34 | - |
| | FALCON-Unzip | 3528 | 30 | - |
| | PECAT | 142 | 41 | 49 |
| *B. taurus* Angus×Brahman, CLR, 135X | PECAT | 4437 | 219 | 1099 |
| *A. thaliana* Col-0×C24, ONT, 106X | Canu | 8359 | 29 | - |
| | Flye+Hapdup | 202 | 94 | - |
| | Shasta | 9 | 60 | - |
| | PECAT | 359 | 179 | 142 |
| *B. taurus* Bison×Simmental, ONT, 200X | Flye+Hapdup | 8159 | 387 | - |
| | Shasta | 519 | 1357 | - |
| | PECAT | 8869 | 381 | 1574 |
| HG002, ONT, R9, 59X | Flye+Hapdup | 3020 | 430 | - |
| | Shasta | 153 | 781 | - |
| | PECAT | 7456 | 348 | 1211 |

All the pipelines are tested on the same computer with a 2.0 GHz CPU and 2T GB RAM of memory and run with 48 threads. Since FALCON-Unzip and Canu didn't generate the assemblies in 3 weeks, it is excluded on datasets *B. taurus* and HG002.

**Supplementary Table 12. Performance comparison of error correction by PECAT on HG002 datasets using different sequencing techniques.**

| Dataset | Pipeline | N50 | Match (%) | Mismatch (%) | Insertion (%) | Deletion (%) | Consistency (%) | Completeness PAT (%) / MAT (%) |
|---|---|---|---|---|---|---|---|---|
| ONT, R9, UL, 59X | raw data | 107537 | 94.97 | 1.81 | 1.24 | 1.97 | 98.75 | 98.6 / 98.6 |
| | PECAT | 104009 | 99.85 | 0.04 | 0.04 | 0.08 | 99.85 | 98.8 / 99.1 |
| ONT, R10, UL, 116X | raw data | 235027 | 98.25 | 0.61 | 0.48 | 0.66 | 99.64 | 99.4 / 99.7 |
| | PECAT | 210995 | 99.88 | 0.02 | 0.03 | 0.06 | 99.98 | 99.3 / 99.7 |
| ONT, R10, Duplex, 45X | raw data | 37531 | 99.67 | 0.08 | 0.09 | 0.17 | 99.80 | 99.4 / 99.8 |
| | PECAT | 36780 | 99.96 | 0.00 | 0.01 | 0.03 | 99.96 | 99.2 / 99.6 |
| HiFi, 36X | raw data | 14718 | 99.77 | 0.01 | 0.10 | 0.11 | 99.87 | 99.4 / 99.8 |
| | PECAT | 14661 | 99.99 | 0.00 | 0.00 | 0.01 | 99.98 | 99.2 / 99.6 |

'Match', 'Mismatch', 'Insertion', and 'Deletion' are the percentages of matches, mismatches, insertions, and deletions in the alignments between the reads and the reference genome. 'Size' is the total number of base pairs of the corrected reads. 'N50' is the length of the shortest read for which longer and equal-length reads cover at least 50% of the 40X longest reads. 'Consistency' is defined as $\sum\max(k_p, k_m)/\sum(k_p + k_m)$, in which $k_p$ and $k_m$ are the number of paternal and maternal haplotype-specific k-mers in each read. 'Completeness' is the percentage of parent-specific k-mers (occurrences $\geq 4$) in the 40X longest reads. 'PAT' and 'MAT' are paternal Completeness and maternal Completeness respectively. 'ONT' indicates the dataset is composed of Nanopore reads. 'UL' indicates the reads are ultra-long reads. 'Duplex' indicates the dataset is generated by the duplex sequencing method. 'HiFi' indicates the dataset is composed of PacBio HiFi reads.

**Supplementary Table 13. Precision, recall, and F1-score of small variants (SNP, INDEL) and structural variants (SV) in HG002 assemblies from Nanopore R10 (ultra-long) reads.**

| Pipeline | SNP | | | INDEL | | | SV | | |
|---|---|---|---|---|---|---|---|---|---|
| | Precision (%) | Recall (%) | F1-score (%) | Precision (%) | Recall (%) | F1-score (%) | Precision (%) | Recall (%) | F1-score (%) |
| Flye+Hapdup (dual) | 99.6 | 99.5 | 99.6 | 64.5 | 84.3 | 73.1 | 94.7 | 97.8 | 96.2 |
| Shasta (pri/alt) | 99.4 | 98.7 | 99.0 | 63.0 | 72.5 | 67.4 | 94.9 | 94.4 | 94.7 |
| PECAT (pri/alt) | 99.5 | 99.3 | 99.4 | 71.6 | 87.6 | 78.8 | 94.7 | 98.0 | 96.3 |
| PECAT (dual) | 99.5 | 99.5 | 99.5 | 71.6 | 87.8 | 78.9 | 94.7 | 98.0 | 96.3 |

‘pri/alt’ represents primary/alternate format. ‘dual’ represents dual assembly format.

**Supplementary Table 14. Performance of assemblies by PECAT on HG002 Nanopore datasets with different coverage.**

| Dataset | Format | Size (Mb) | NG50 (Mb) | Quality (reference-based) | Quality (k-mer-based) | BUSCO (%) | Hamming error (%) | Phase block NG50 (Mb) | Intra-block switch error (%) |
|---|---|---|---|---|---|---|---|---|---|
| Ref | - | 2959.3/3061.7 | 146.7/154.4 | -/- | 58.6/59.4 | 92.8/95.8 | 0.15/0.08 | 90.4/106.7 | 0.02/0.03 |
| HG002 ONT, 37X | pri/alt | 3061.4/2772.9 | 74.8/7.7 | 30.3/30.3 | 39.4/39.3 | 94.1/89.5 | 18.84/0.89 | 11.8/6.7 | 0.09/0.12 |
| | dual | 3055.7/2829.5 | 60.5/59.3 | 30.3/30.4 | 39.4/40.1 | 94.0/91.1 | 12.05/15.43 | 17.7/17.1 | 0.09/0.13 |
| HG002 ONT, 59X | pri/alt | 3059.3/2857.7 | 92.9/15.0 | 30.8/30.8 | 41.8/41.6 | 94.7/91.0 | 15.72/1.48 | 22.2/13.0 | 0.08/0.11 |
| | dual | 3057.2/2927.2 | 92.7/74.5 | 30.8/30.8 | 41.8/41.8 | 94.6/91.6 | 9.67/10.98 | 30.8/23.6 | 0.08/0.11 |

‘Size’ is the total number of base pairs in all contigs generated by assemblers. ‘NG50’ is the length of the shortest contig for which longer and equal length contigs cover at least 50 of genome size. The genome size of HG002 that we used for evaluation is 3G, respectively. ‘BUSCO’ is gene completeness evaluated by BUSCO. ‘Quality (reference-based)’ is the metric ‘q50’ evaluated by Pomoxis. ‘Hamming error’ is the fraction of nondominant parental-specific k-mers in a contig. ‘Quality (k-mer-based)’, ‘Phase block NG50’, and ‘Intra-block switch error’ are evaluated by mercury. ‘pri/alt’ represents primary/alternate assembly format. ‘dual’ represents dual assembly format. The two sets of contigs are separately reported in each cell. ‘Ref’ is the reference genome. The sources of the reference genomes are illustrated in **Supplementary Table 18**.

**Supplementary Table 15. Performance comparison of assembly on Nanopore R10 and PacBio HiFi reads.**

| Dataset | Pipeline | Size (Mb) | NG50 (Mb) | Quality (reference-based) | Quality (k-mer-based) | BUSCO (%) | Hamming error (%) | Phase block NG50 (Mb) | Intra-block switch error (%) |
|---|---|---|---|---|---|---|---|---|---|
| | Ref | 2959.3/3061.7 | 146.7/154.4 | -/- | 58.6/59.4 | 92.8/95.8 | 0.15/0.08 | 90.4/106.7 | 0.02/0.03 |
| HG002 ONT, R10, UL, 116X | Flye+HapDup (dual) | 2954.7/2952.1 | 61.9/59.3 | 36.4/36.2 | 48.9/48.2 | 95.8/95.7 | 3.13/3.02 | 46.5/39.1 | 0.02/0.03 |
| | Shasta (pri/alt) | 3095.4/2729.9 | 45.3/33.8 | 34.3/34.4 | 42.1/48.7 | 95.7/92.2 | 3.05/0.86 | 17.2/15.4 | 0.30/0.44 |
| | PECAT (pri/alt) | 3159.7/2895.6 | 91.4/59.9 | 38.0/38.0 | 49.0/50.1 | 95.8/92.4 | 3.75/0.30 | 59.4/58.0 | 0.04/0.05 |
| | PECAT (dual) | 3153.4/2917.8 | 91.4/80.2 | 38.0/38.0 | 49.2/50.0 | 95.8/92.7 | 2.99/1.49 | 63.8/59.2 | 0.04/0.06 |
| HG002 ONT, R10, Duplex, 45X | Flye+HapDup (dual) | 2921.2/2921.7 | 26.1/26.1 | 37.2/37.2 | 52.2/51.9 | 95.7/95.8 | 20.61/20.90 | 4.2/4.2 | 0.04/0.04 |
| | Shasta (pri/alt) | 3116.3/2292.6 | 38.6/0.8 | 36.8/36.0 | 48.6/57.0 | 95.4/75.3 | 21.11/0.31 | 2.1/0.7 | 0.28/0.23 |
| | PECAT (pri/alt) | 3035.8/2762.6 | 75.6/1.1 | 38.5/38.3 | 55.1/54.3 | 95.8/86.7 | 24.55/0.42 | 2.6/0.9 | 0.07/0.08 |
| | PECAT (dual) | 3029.0/2828.0 | 71.2/70.8 | 38.5/38.5 | 55.5/55.9 | 95.6/92.6 | 22.48/34.47 | 4.3/3.7 | 0.07/0.10 |
| HG002 HiFi, 36X | Flye+HapDup (dual) | 3211.4/3210.9 | 4.6/4.6 | 47.0/45.2 | 53.4/52.8 | 94.7/94.7 | 14.88/14.93 | 0.7/0.7 | 0.21/0.22 |
| | Hifiasm (pri/alt) * | 3112.2/2910.3 | 89.9/0.4 | 50.0/50.0 | 55.3/56.5 | 95.6/77.0 | 24.70/0.37 | 1.1/0.3 | 0.11/0.01 |
| | Hifiasm (dual) * | 3015.3/3077.5 | 44.8/64.5 | 50.0/50.0 | 54.9/57.8 | 95.5/94.9 | 34.80/24.10 | 1.0/1.0 | 0.16/0.11 |
| | PECAT (pri/alt) | 2991.8/2344.4 | 50.9/0.2 | 45.2/46.0 | 53.3/52.3 | 95.6/67.4 | 25.21/0.32 | 0.7/0.1 | 0.19/0.07 |
| | PECAT (dual) | 2990.8/2798.7 | 41.0/39.7 | 45.2/45.2 | 53.4/57.1 | 95.6/92.3 | 24.02/35.49 | 0.9/0.7 | 0.19/0.27 |

'Size' is the total number of base pairs in all contigs generated by assemblers. 'NG50' is the length of the shortest contig for which longer and equal length contigs cover at least 50 of genome size. The genome sizes of HG002 that we used for evaluation is 3G. 'BUSCO' is gene completeness evaluated by BUSCO. 'Hamming error' is the fraction of nondominant parental-specific k-mers in a contig. 'Quality (reference-based)' is the metric 'q50' evaluated by Pomoxis. 'Quality (k-mer-based)', 'Phase block NG50', and 'Intra-block switch error' are evaluated by mercury. 'pri/alt' represents primary/alternate assembly format. 'dual' represents dual assembly format. 'ONT' indicates the dataset is composed of Nanopore reads. 'UL' indicates the reads are ultra-long reads. 'Duplex' indicates the dataset is generated by the duplex sequencing method. 'HiFi' indicates the dataset is composed of PacBio HiFi reads. The two sets of contigs are separately reported in each cell. 'Ref' is the reference genome. The sources of the reference genomes are illustrated in **Supplementary Table 18**. Asterisks mark previously published assemblies.

**Supplementary Table 16. Precision, recall, and F1-score of small variants (SNP, INDEL) and structural variants (SV) in HG002 assemblies from Nanopore R10 duplex reads.**

| Pipeline | SNP | | | INDEL | | | SV | | |
|---|---|---|---|---|---|---|---|---|---|
| | Precision (%) | Recall (%) | F1-score (%) | Precision (%) | Recall (%) | F1-score (%) | Precision (%) | Recall (%) | F1-score (%) |
| Flye+Hapdup (dual) | 99.7 | 99.5 | 99.6 | 64.9 | 85.3 | 73.8 | 94.6 | 97.9 | 96.2 |
| Shasta (pri/alt) | 99.7 | 89.7 | 94.4 | 70.1 | 66.0 | 68.0 | 94.8 | 96.2 | 95.5 |
| PECAT (pri/alt) | 99.6 | 96.9 | 98.2 | 74.5 | 85.9 | 79.8 | 93.7 | 97.9 | 95.8 |
| PECAT (dual) | 99.6 | 99.3 | 99.4 | 74.3 | 88.5 | 80.8 | 94.5 | 97.9 | 96.2 |

‘pri/alt’ represents primary/alternate format. ‘dual’ represents dual assembly format.

**Supplementary Table 17. Precision, recall, and F1-score of small variants (SNP, INDEL) and structural variants (SV) in HG002 assemblies from HiFi reads.**

| Pipeline | SNP | | | INDEL | | | SV | | |
|---|---|---|---|---|---|---|---|---|---|
| | Precision (%) | Recall (%) | F1-score (%) | Precision (%) | Recall (%) | F1-score (%) | Precision (%) | Recall (%) | F1-score (%) |
| Flye+Hapdup (dual) | 99.4 | 95.6 | 97.5 | 96.3 | 95.7 | 96.0 | 94.5 | 93.4 | 94.0 |
| Hifiasm (pri/alt) | 99.8 | 92.4 | 96.0 | 97.9 | 90.4 | 94.0 | 92.6 | 98.1 | 95.3 |
| Hifiasm (dual) | 99.6 | 99.0 | 99.3 | 97.4 | 97.1 | 97.3 | 94.5 | 97.8 | 96.1 |
| PECAT (pri/alt) | 99.7 | 85.0 | 91.8 | 96.2 | 82.7 | 88.9 | 93.8 | 97.5 | 95.6 |
| PECAT (dual) | 99.5 | 98.7 | 99.1 | 96.5 | 96.8 | 96.6 | 94.8 | 97.5 | 96.1 |

‘pri/alt’ represents primary/alternate format. ‘dual’ represents dual assembly format.

**Supplementary Table 18. Detailed information of the datasets used in this study.**

| Dataset | Type | Data resource |
|---|---|---|
| NCTC9024<br>NCTC9007 | PacBio CLR | http://www.sanger.ac.uk/resources/downloads/bacteria/nctc/ |
| *S. cerevisiae*<br>SK1×Y12 | PacBio CLR | ERR1080522, ERR1080529, ERR1080536, ERR1080537, ERR1124245, ERR1140978 (SK1)<br>ERR1080526, ERR1080538, ERR1080539, ERR1140975, ERR1140979, ERR985361 (Y12) |
| | NGS | SRR4074258(SK1)<br>SRR4074358(Y12) |
| | Reference | ASM205788v1 (SK1)<br>ASM205864v1 (Y12) |
| *A. thaliana*<br>Col-0×Cvi-0 | PacBio CLR | SRR3405242 - SRR3405290 (Col-0)<br>SRR3405327 - SRR3405386 (Cvi-0)<br>SRR3405291 - SRR3405326 (F1) |
| | NGS | SRR3703081, SRR3703082, SRR3703105 (F1)<br>SRR16841689 (Col-0)<br>ERR3624578 (Cvi-0) |
| | Reference | GCA_020911765.1 (Col-0)<br>GCA_902460275.1 (Cvi-0) |
| *D. melanogaster*<br>ISO1×A4 | PacBio CLR | SRR9969843 (F1) |
| | NGS | SRR6702604 (ISO1)<br>SRR457665, SRR457666, SRR457707 (A4) |
| | Reference | GCA_000001215.4 (ISO1)<br>ASM340174v1 (A4) |
| *B. taurus*<br>Angus ×<br>Brahman | PacBio CLR | SRR6691718, SRR6691728 - SRR6691879, SRR6691882 -SRR6691900, SRR6691904, SRR6691905, SRR6691908 -SRR6691950, SRR6691954 - SRR6691960, SRR6691962 - SRR6691984 (F1) |
| | NGS | SRR6691901 - SRR6691903, SRR6691907 (Angus)<br>SRR6691719, SRR6691880, SRR6691881, SRR6691906 (Brahman)<br>SRR6691720 - SRR6691727, SRR6691748,<br>SRR6691951- SRR6691953, SRR6691961 (F1) |
| | Reference | GCA_003369685.2 (Angus)<br>GCA_003369695.2 (Brahman) |
| *A. thaliana*<br>Col-0×C24 | Nanopore | CRA008108 (F1) * |
| | NGS | SRR16841689 (Col-0)<br>ERR3624577 (C24) |
| | Reference | GCA_020911765.1 (Col-0)<br>GCA_902705455.1 (C24) |
| *B. taurus*<br>Bison ×<br>Simmental | Nanopore | SRR13105460 - SRR13105478 (F1) |
| | NGS | SRR13081814 (Bison)<br>SRR13081923 (Simmental)<br>SRR13081717 (F1) |

| | | |
|---|---|---|
| | Reference | GCA_018282365.1 (Bison)<br>GCA_018282465.1 (Simmental) |
| HG002 | Nanopore R9 | https://s3-us-west-2.amazonaws.com/human-pangenomics/index.html?prefix=T2T/scratch/HG002/sequencing/ont/<br>03_08_22_R941_HG002_1_Guppy_6.0.6_prom_sup.fastq.gz<br>03_08_22_R941_HG002_2_Guppy_6.0.6_prom_sup.fastq.gz |
| | Nanopore R10 | https://s3-us-west-2.amazonaws.com/human-pangenomics/index.html?prefix=submissions/5b73fa0e-658a-4248-b2b8-cd16155bc157--UCSC_GIAB_R1041_nanopore/HG002_R1041_UL/Guppy6/ |
| | Nanopore R10 Duplex | https://s3-us-west-2.amazonaws.com/human-pangenomics/index.html?prefix=submissions/0CB931D5-AE0C-4187-8BD8-B3A9C9BFDADE--UCSC_HG002_R1041_Duplex_Dorado/Dorado_v0.1.1/stereo_duplex |
| | PacBio HiFi | SRR10382244, SRR10382245, SRR10382248, SRR10382249 |
| | NGS | https://s3-us-west-2.amazonaws.com/human-pangenomics/index.html?prefix=NHGRI_UCSC_panel/HG002/hpp_HG002_NA24385_son_v1/ |
| | Reference | GCA_021950905.1 (HG003)<br>GCA_021951015.1 (HG004) |

NGS: Next-generation sequencing. The datasets marked with asterisks are available at https://ngdc.cncb.ac.cn/. The other datasets are available at https://www.ncbi.nlm.nih.gov/.
