## Supplementary Figures for "*de novo* diploid genome assembly using long noisy reads"

a

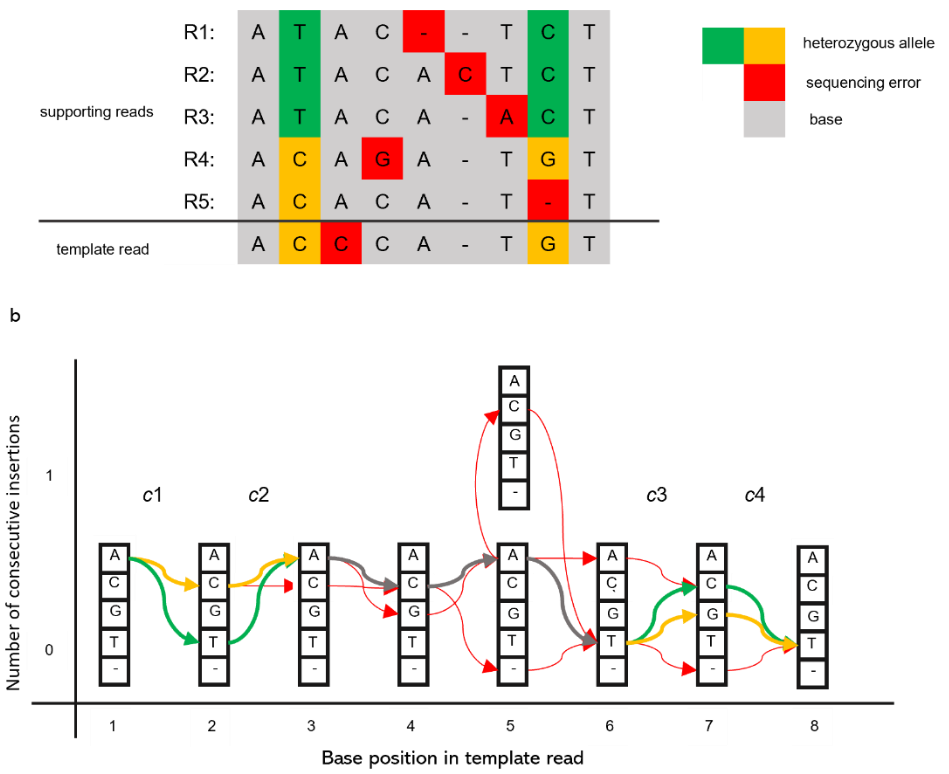


c

| Read | c1 | c2 | c3 | c4 | Total |
|---|---|---|---|---|---|
| R1: | -1 | 0 | -1 | -1 | -3 |
| R2: | -1 | 0 | -1 | -1 | -3 |
| R3: | -1 | 0 | 0 | -1 | -2 |
| R4: | 1 | 0 | 1 | 1 | 3 |
| R5: | 1 | 0 | 0 | 0 | 1 |

d

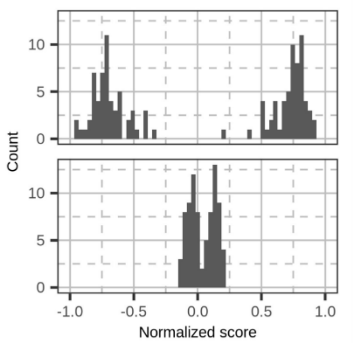


**Supplementary Figure 1. Illustration of haplotype-aware error correction.** (**a**) The alignments between the template read and the five supporting reads R1-5. The heterozygous alleles and sequencing errors are marked with different colors. (**b**) The POA graph generated from the alignments. The red edges are supported by only one read, and they are considered sequencing errors. Other thick edges are supported by multiple reads. The edges passing through the heterozygous alleles are marked with corresponding colors. There four important locations *c*1-4 are identified in the POA graph, at which there are two dominant edges. (**c**) The scores of the supporting reads at important locations. If the supporting read and the template read pass through the same dominant edge, the score is 1 and if they pass through the different dominant edges, the score is -1. If one of them doesn't pass through any dominant edge, which means there is a sequencing error, the score is 0. (**d**) The histograms of normalized scores of two supporting reads from the HG002 dataset. Two peaks indicate that the supporting reads are from two different haplotypes. PECAT selects the reads whose scores fall into the first peak for error correction.

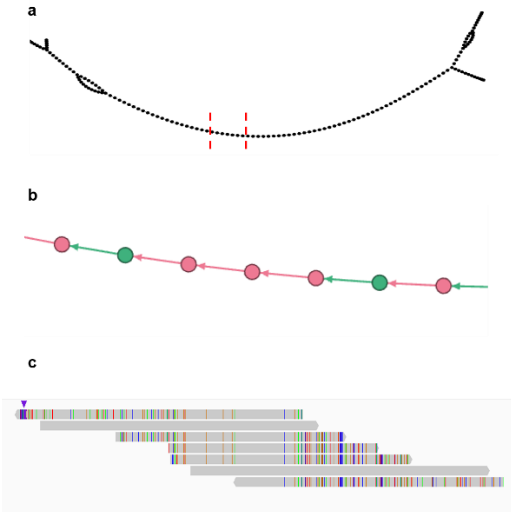


**Supplementary Figure 2. Inconsistent overlaps in the string graph**. (**a**) Part of the string graph built from *S. cerevisiae* (SK1 × Y12) corrected reads. (**b**) Enlarged view of the red dashed area in **a**. Each node represents a read and is colored according to its haplotype. (**c**) The overlaps between the reads in **b**. Colored dashes indicate haplotype differences.

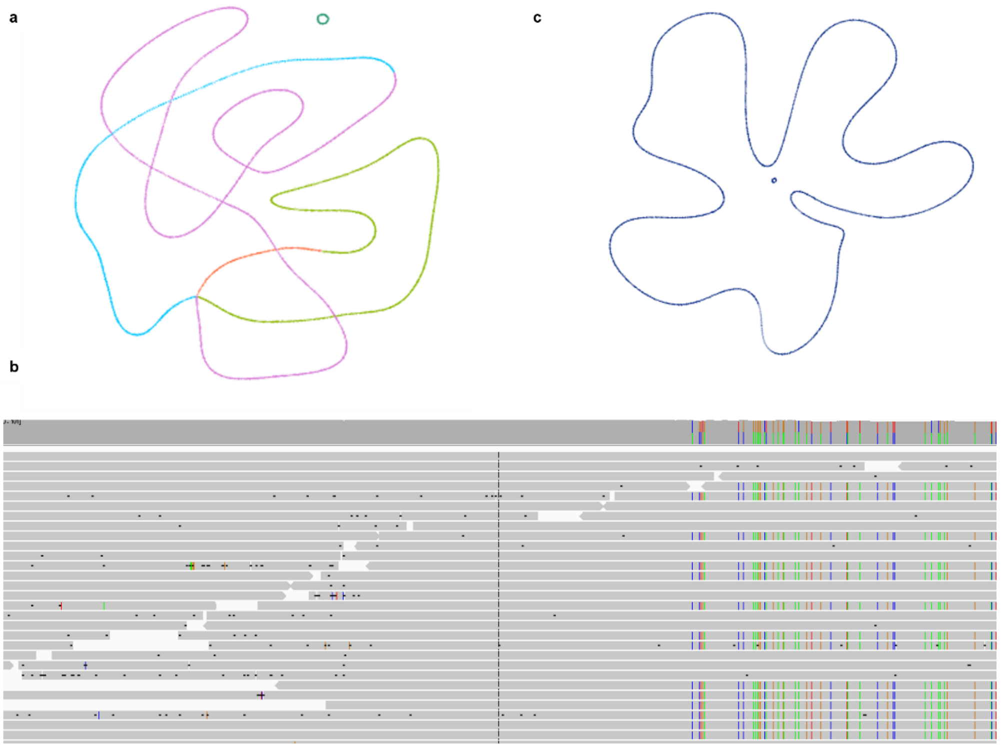


**Supplementary Figure 3. Illustration of PECAT resolving the repeat of NCTC9006.** (**a**) The assembly graph generated by PECAT in the first round of assembly. Different colors represent different contigs. There is an unresolvable double repeat in the graph. (**b**) The alignment between the corrected reads and the repeat. Colored dashed indicate differences. PECAT identifies the inconsistent overlaps according to the differences. (**c**) The assembly graph generated by PECAT in the second round of assembly. PECAT resolves the repeat by filtering out inconsistent overlaps in the second round of assembly.

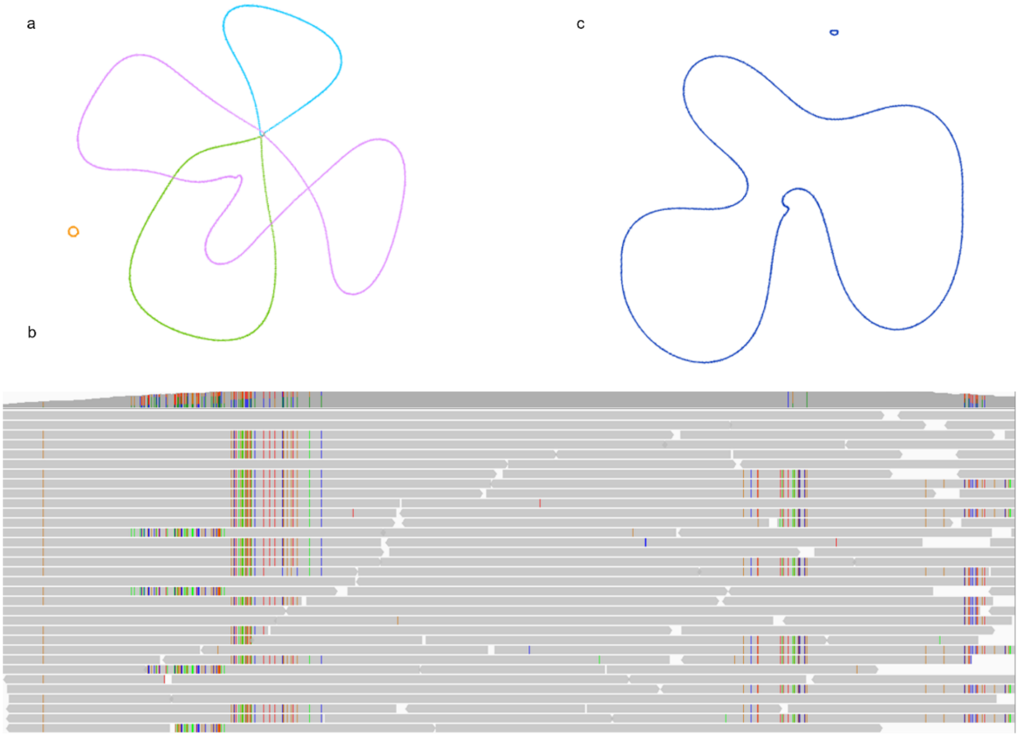


**Supplementary Figure 4. Illustration of PECAT resolving the repeat of NCTC9024.** (**a**) The assembly graph generated by PECAT in the first round of assembly. Different colors represent different contigs. There is an unresolvable triple repeat in the graph. (**b**) The alignments between the corrected reads and the repeat. Colored dashed indicate differences. PECAT identifies the inconsistent overlaps according to the differences. (**c**) The assembly graph generated by PECAT in the second round of assembly. PECAT resolves the repeat by filtering out inconsistent overlaps in the second round of assembly.

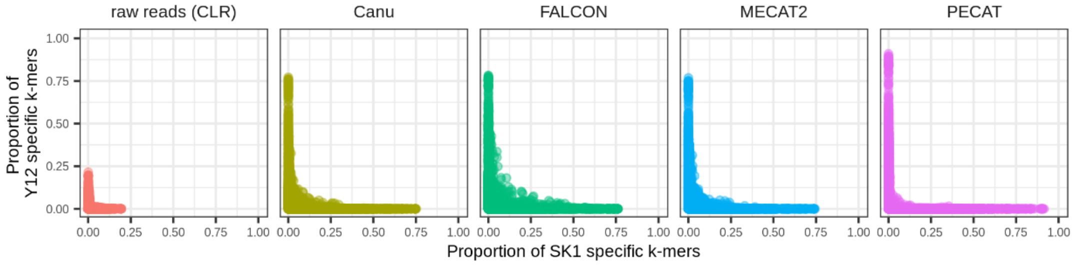


**Supplementary Figure 5. Haplotype-specific k-mers consistency of *S. cerevisiae* (SK1 × Y12) raw reads and corrected reads by different methods.** Each point corresponds to a read. Its coordinate gives the proportion of the parental specific k-mers in the read, where k is 17. All 40X longest reads are shown in each subfigure.

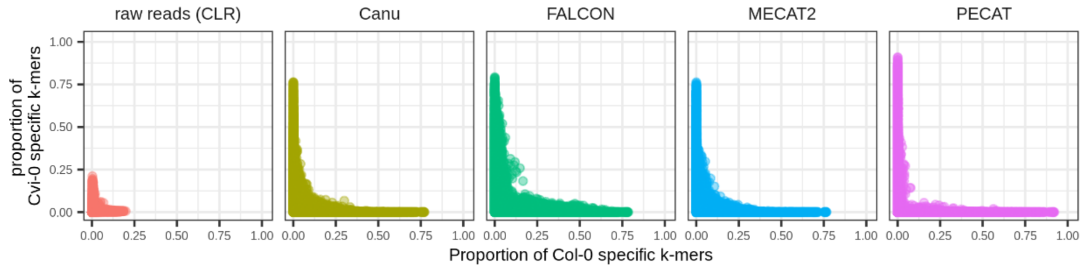


**Supplementary Figure 6. Haplotype-specific k-mers consistency of *A. thaliana* (Col-0 × Cvi-0) raw reads and corrected reads by different methods.** Each point corresponds to a read. Its coordinate gives the proportion of the parental specific k-mers in the read, where k is 18. All 40X longest reads are shown in each subfigure.

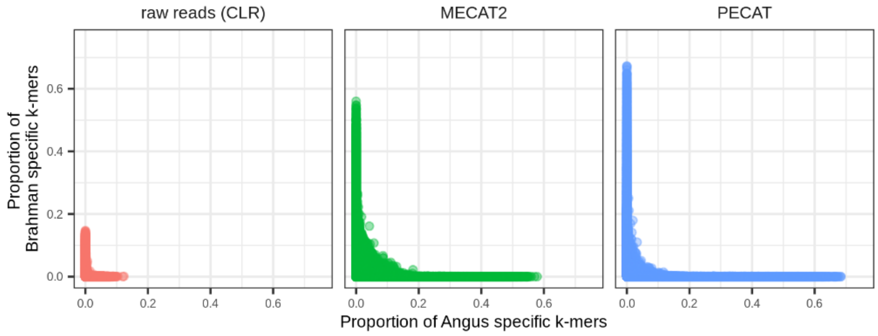


**Supplementary Figure 7. Haplotype-specific k-mers consistency of *B. taurus* (Angus×Brahman) raw reads and corrected reads by different methods.** Each point corresponds to a read. Its coordinate gives the proportion of the parental specific k-mers in the read, where k is 21. All 40X longest reads are shown in each subfigure.

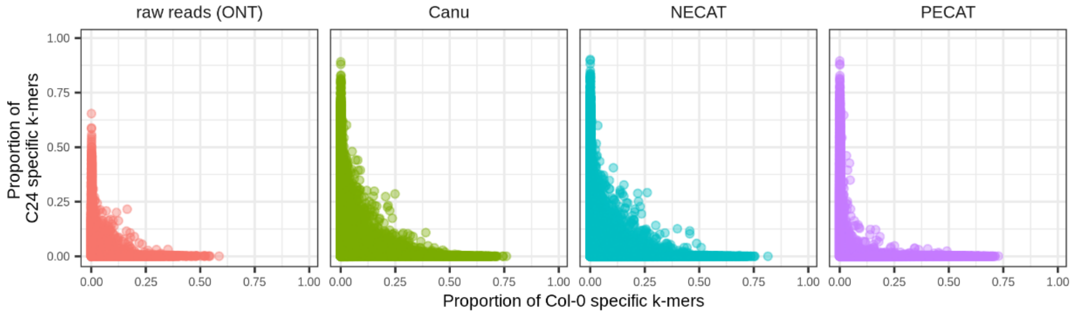


**Supplementary Figure 8. Haplotype-specific k-mers consistency of *A. thaliana* (Col-0×C24) raw reads and corrected reads by different methods.** Each point corresponds to a read. Its coordinate gives the proportion of the parental specific k-mers in the read, where k is 18. All 40X longest reads are shown in each subfigure.

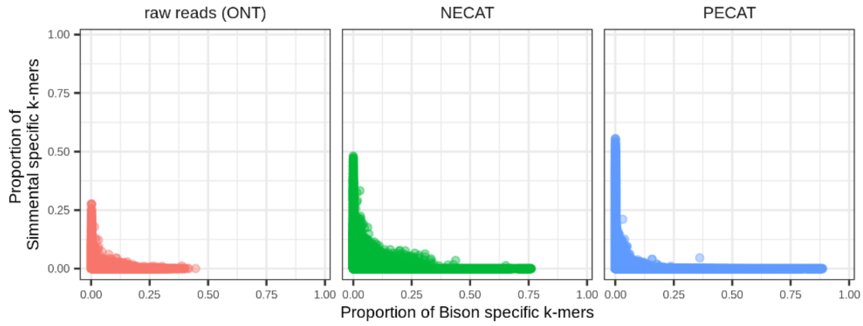


**Supplementary Figure 9. Haplotype-specific k-mers consistency of *B. taurus* (Bison × Simmetal) raw reads and corrected reads by different methods.** Each point corresponds to a read. Its coordinate gives the proportion of the parental specific k-mers in the read, where k is 21. All 40X longest reads are shown in each subfigure.

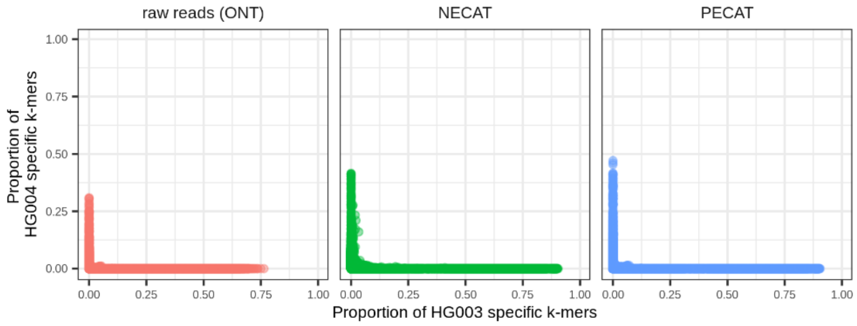


**Supplementary Figure 10. Haplotype-specific k-mers consistency of HG002 R9 raw reads and corrected reads by different methods.** (**a**) Each point corresponds to a read. Its coordinate gives the proportion of the parental specific k-mers in the read, where k is 21. All 40X longest reads are shown in each subfigure.

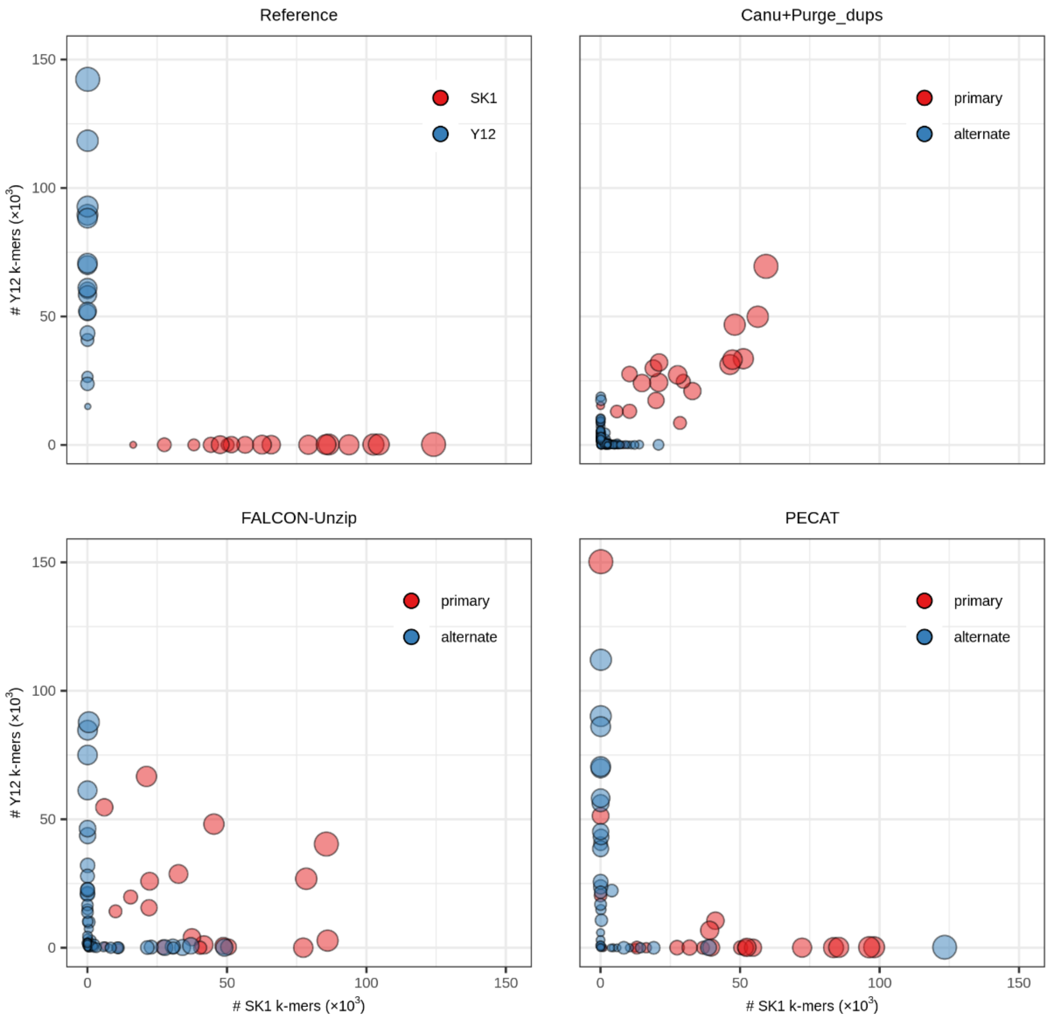


**Supplementary Figure 11. Haplotype-specific k-mer blob plots of the *S. cerevisiae* (SK1 × Y12) reference genome and assemblies by different methods.** All assemblies are in the primary/alternate format. Each blob corresponds to a contig. The coordinate of the blob gives the count of the parental specific k-mers in the contig, where k is 17. Blob size is proportional to contig length.

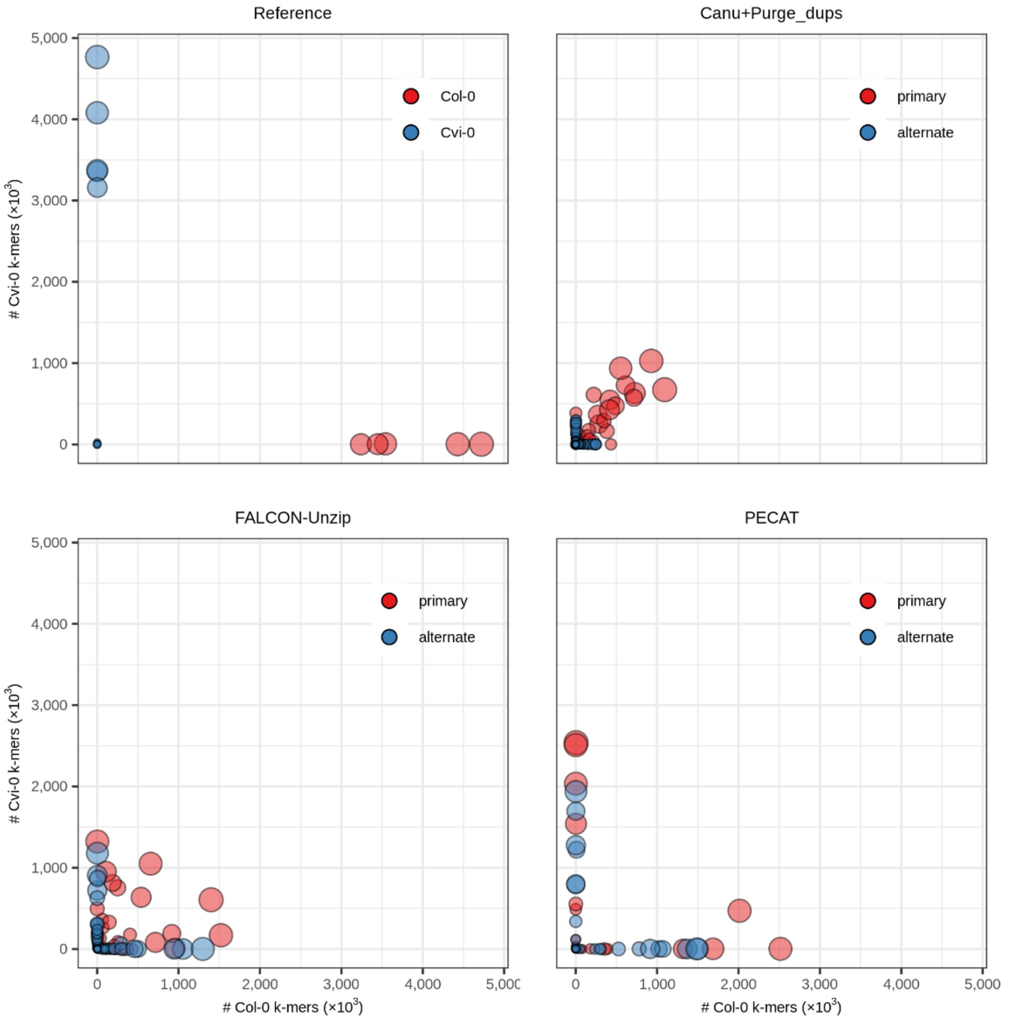


**Supplementary Figure 12. Haplotype-specific k-mer blob plots of the *A. thaliana* (Col-0 × Cvi-0) reference genome and assemblies by different methods.** All assemblies are in the primary/alternate format. Each blob corresponds to a contig. The coordinate of the blob gives the count of the parental specific k-mers in the contig, where k is 18. Blob size is proportional to contig length.

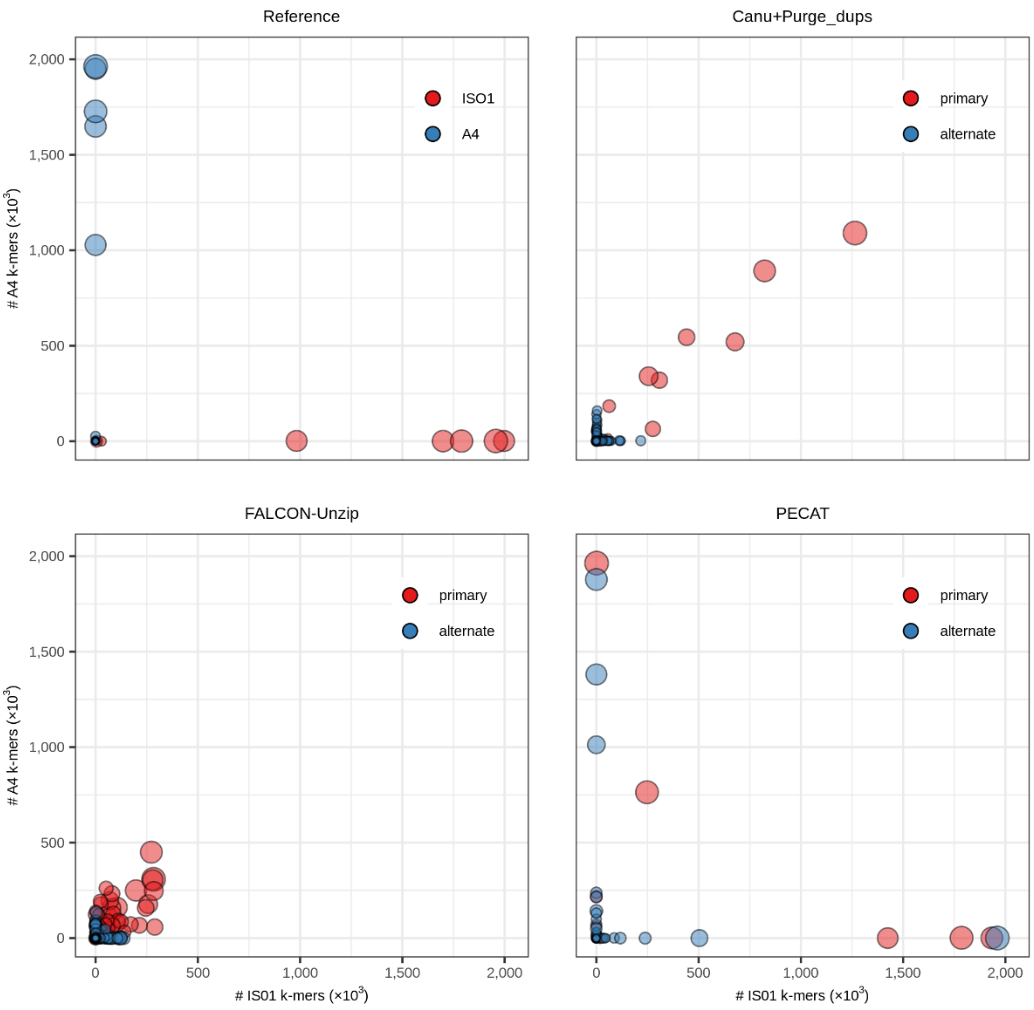


**Supplementary Figure 13. Haplotype-specific k-mer blob plots of the *D. melanogaster* (ISO1 × A4) reference genome and assemblies by different methods.** All assemblies are in the primary/alternate format. Each blob corresponds to a contig. The coordinate of the blob gives the count of the parental specific k-mers in the contig, where k is 18. Blob size is proportional to contig length.

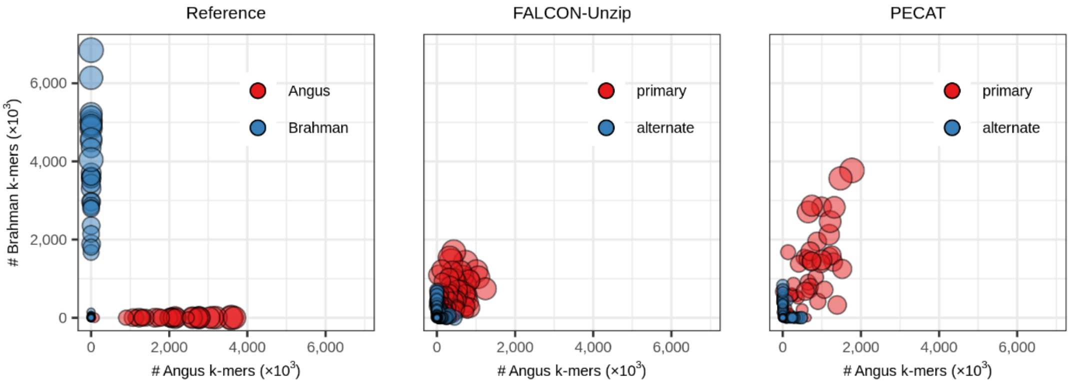


**Supplementary Figure 14. Haplotype-specific k-mer blob plots of the *B. taurus* (Angus×Brahman) reference genome and assemblies by different methods.** All assemblies are in the primary/alternate format. Each blob corresponds to a contig. The coordinate of the blob gives the count of the parental specific k-mers in the contig, where k is 21. Blob size is proportional to contig length.

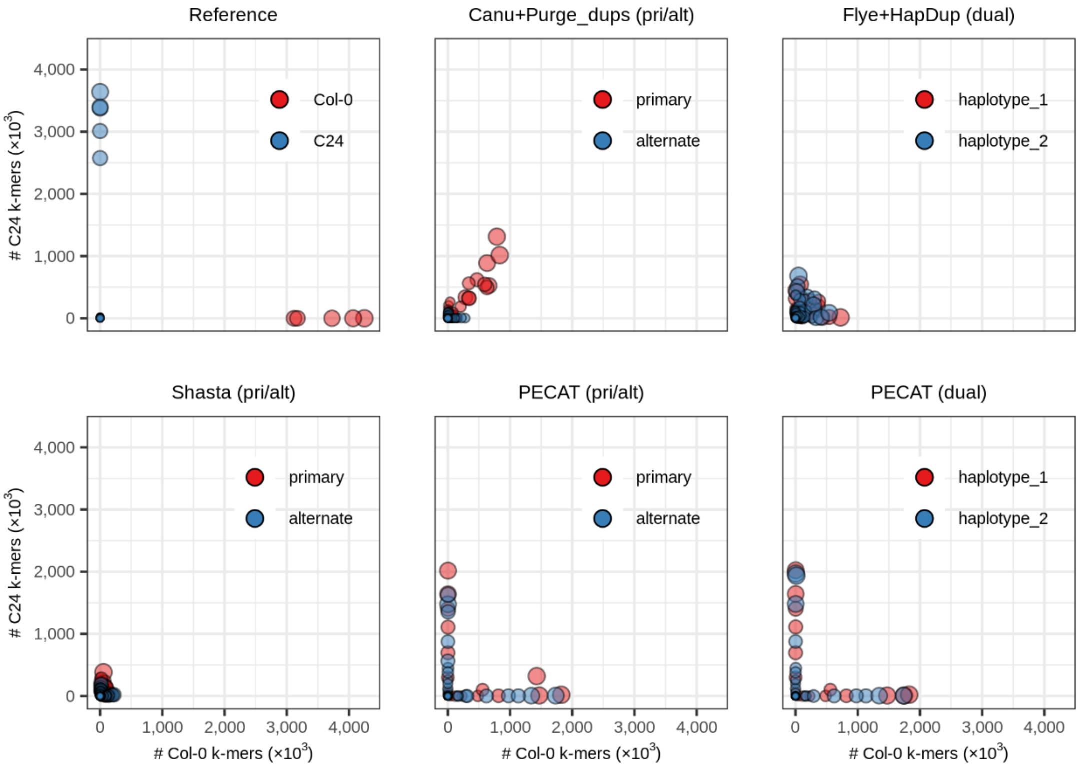


**Supplementary Figure 15. Haplotype-specific k-mer blob plots of the *A. thaliana* (Col-0 × C24) reference genome and assemblies by different methods.** pri/alt or dual represents that the assembly is in the primary/alternate format or the dual assembly format. Each blob corresponds to a contig. The coordinate of the blob gives the count of the parental specific k-mers in the contig, where k is 18. Blob size is proportional to contig length.

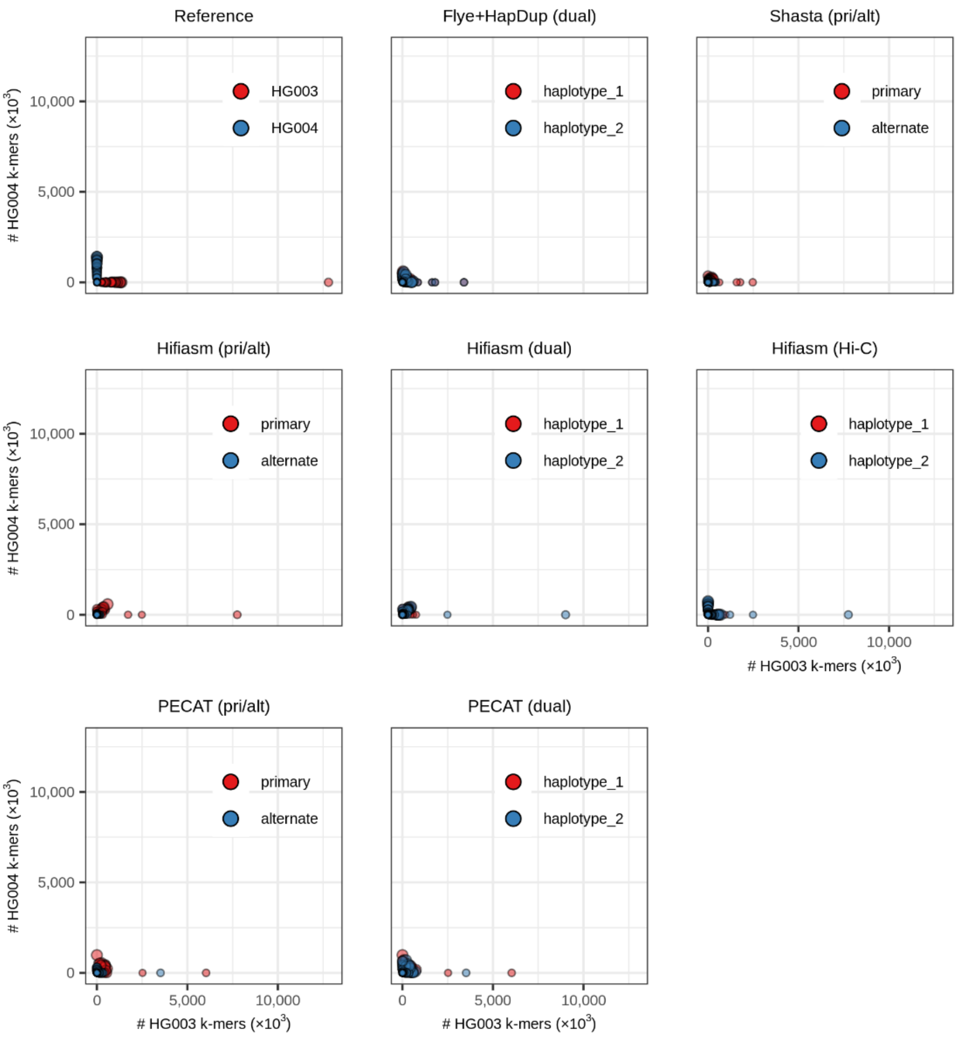


**Supplementary Figure 16. Haplotype-specific k-mer blob plots of the HG002 reference genome and assemblies by different methods.** The assemblies By Hifiasm are from HiFi reads and the others are from Nanopore R9 (ultra-long) reads. Hi-C represents that the assembly uses the additional Hi-C reads. pri/alt or dual represents that the assembly is in the primary/alternate format or the dual assembly format. Each blob corresponds to a contig. The coordinate of the blob gives the count of the parental specific k-mers in the contig, where k is 21. Blob size is proportional to contig length.

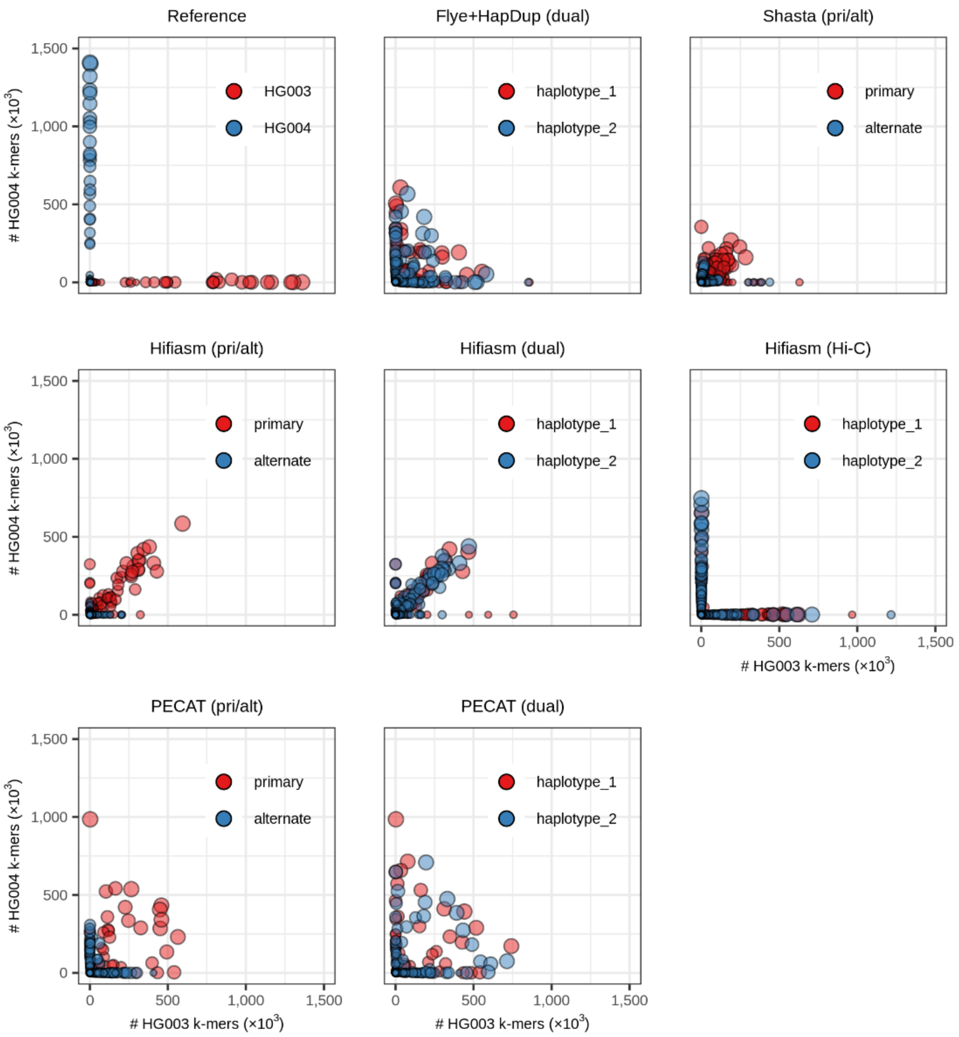


**Supplementary Figure 17**. Enlarged view of **Supplementary Figure 16**.

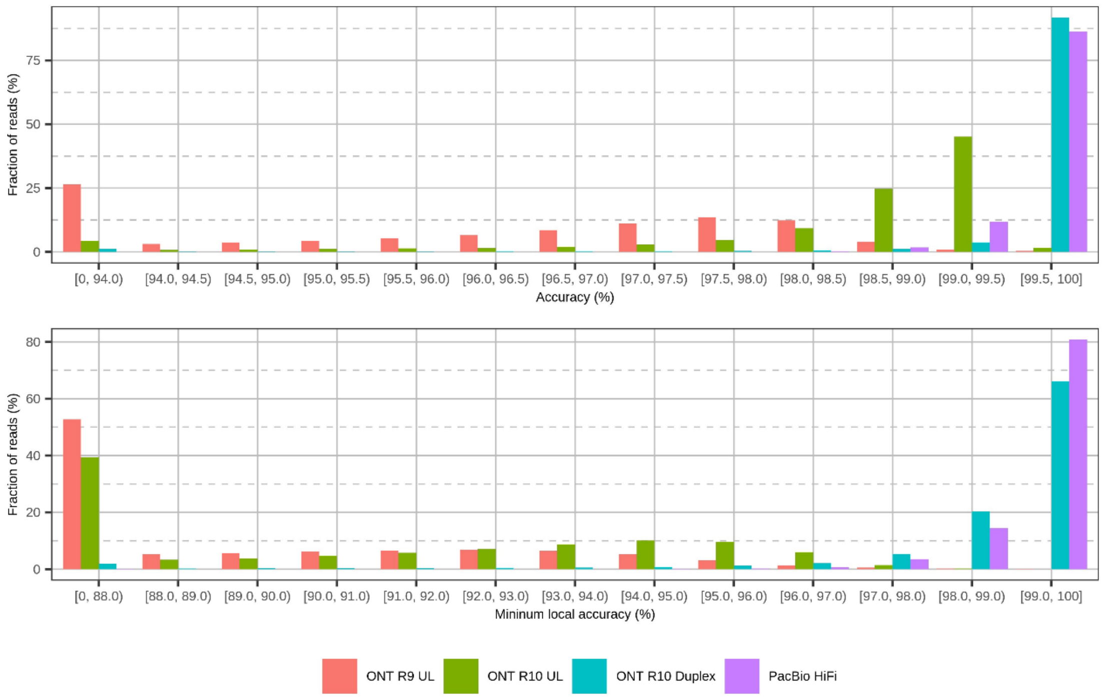


**Supplementary Figure 18. Histogram plot for accuracy of HG002 reads using different sequencing techniques.** ‘Minimum local accuracy’ is the minimum accuracy of windows with 1000 bp in a read. ‘ONT’ indicates the dataset is composed of Nanopore reads. ‘UL’ indicates the reads are ultra-long reads. ‘Duplex’ indicates the dataset is generated by the duplex sequencing method.

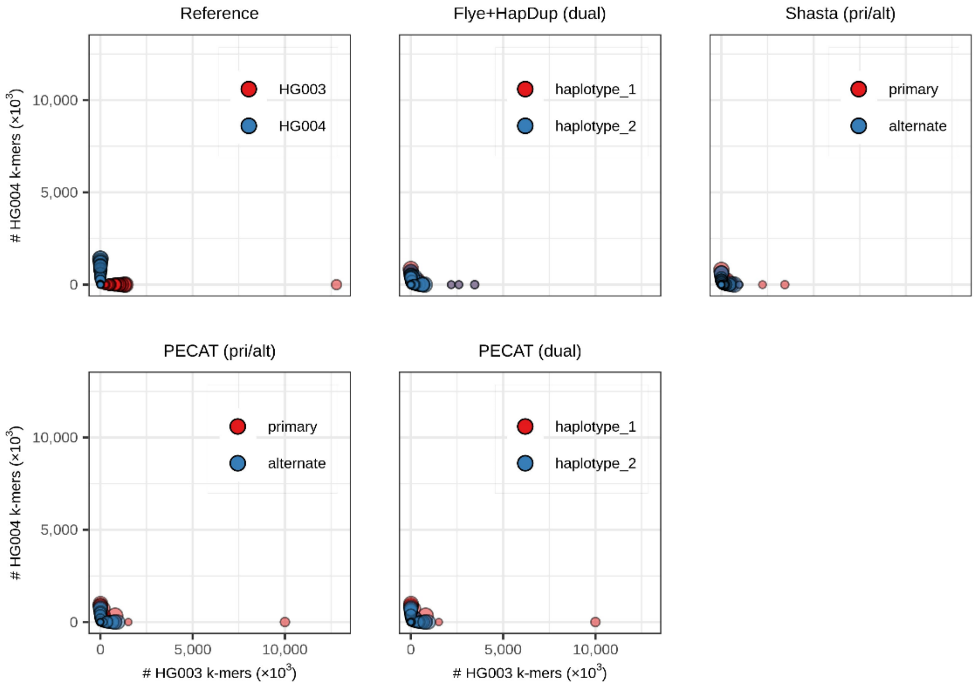


**Supplementary Figure 19. Haplotype-specific k-mer blob plots of the HG002 reference genome and assemblies by different methods from** Nanopore R10 (ultra-long) reads**.** pri/alt or dual represents that the assembly is in the primary/alternate format or the dual assembly format. Each blob corresponds to a contig. The coordinate of the blob gives the count of the parental specific k-mers in the contig, where k is 21. Blob size is proportional to contig length.

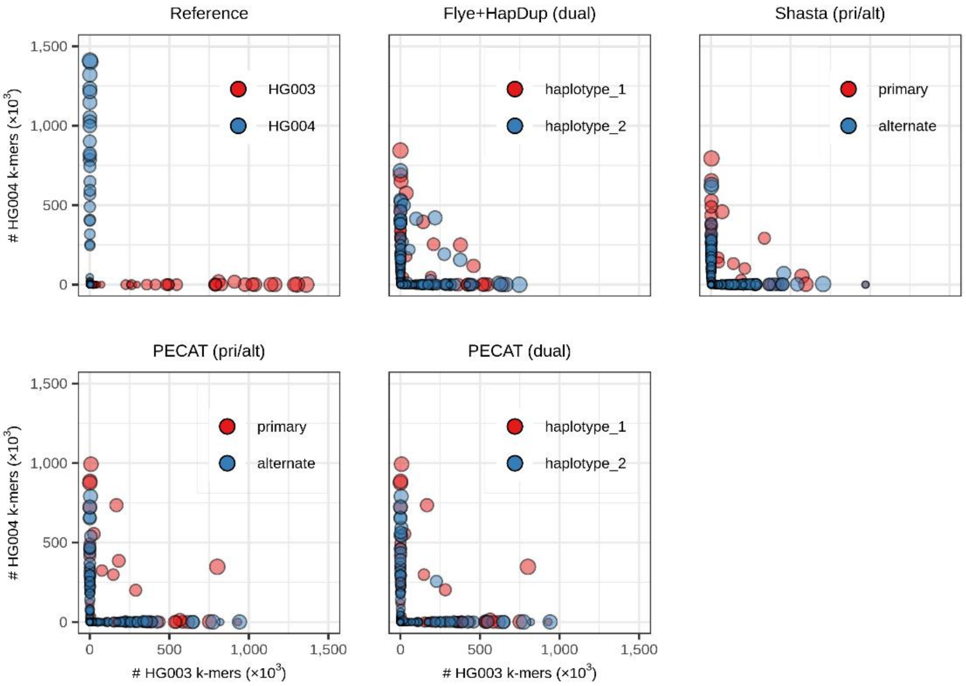


**Supplementary Figure 20**. Enlarged view of **Supplementary Figure 19**.

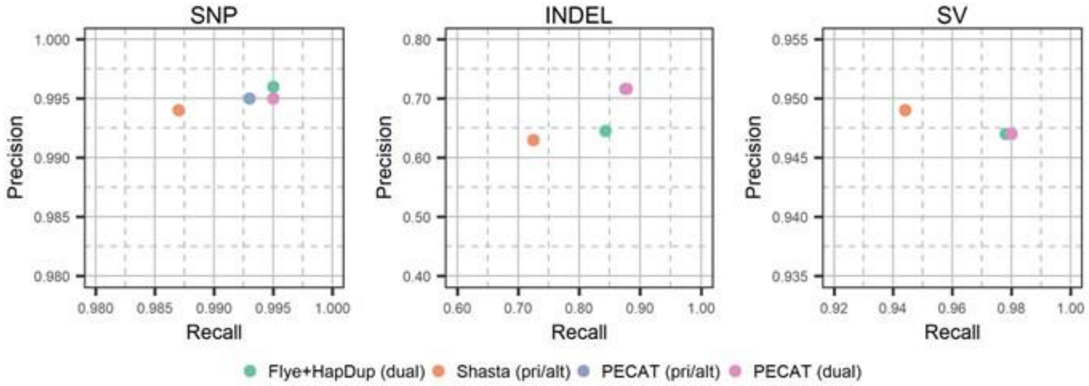


**Supplementary Figure 21**. Precisions and recalls of small variants (SNP, INDEL) and structural variants (SV) in HG002 assemblies from Nanopore R10 (ultra-long) reads. ‘pri/alt’ represents primary/alternate format. ‘dual’ represents dual assembly format. The points of PECAT (pri/alt) and PECAT (dual) overlap in the right two subfigures.

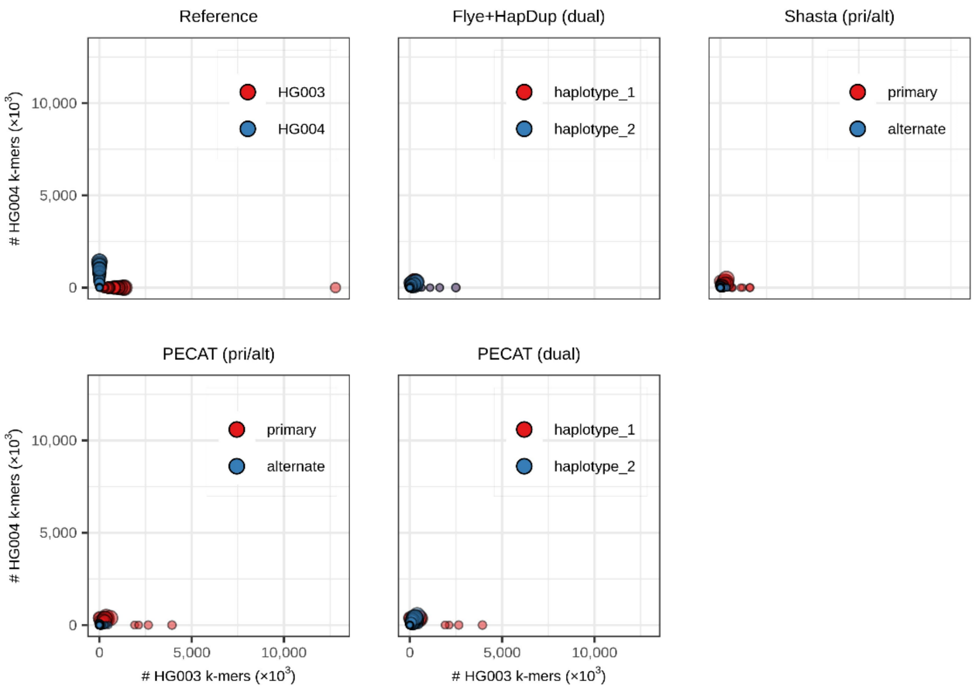


**Supplementary Figure 22. Haplotype-specific k-mer blob plots of the HG002 reference genome and assemblies by different methods from** Nanopore R10 duplex reads**.** pri/alt or dual represents that the assembly is in the primary/alternate format or the dual assembly format. Each blob corresponds to a contig. The coordinate of the blob gives the count of the parental specific k-mers in the contig, where k is 21. Blob size is proportional to contig length.

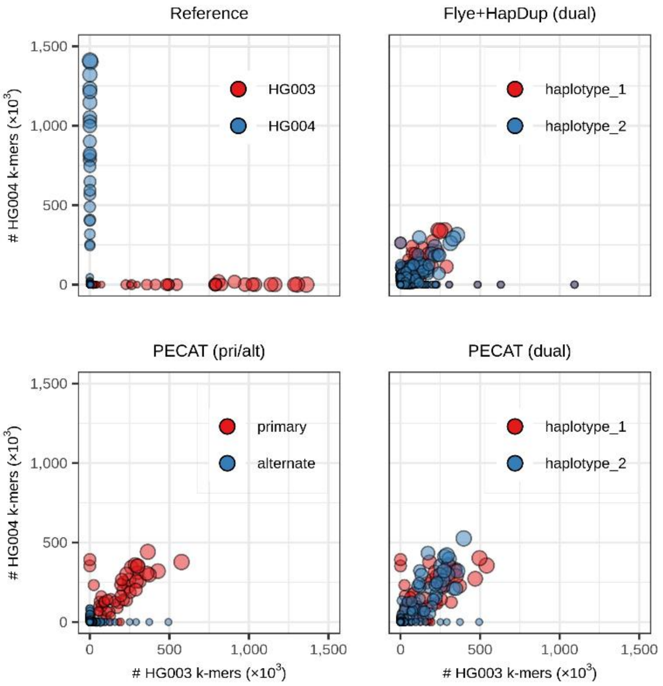


**Supplementary Figure 23**. Enlarged view of **Supplementary Figure 22**.

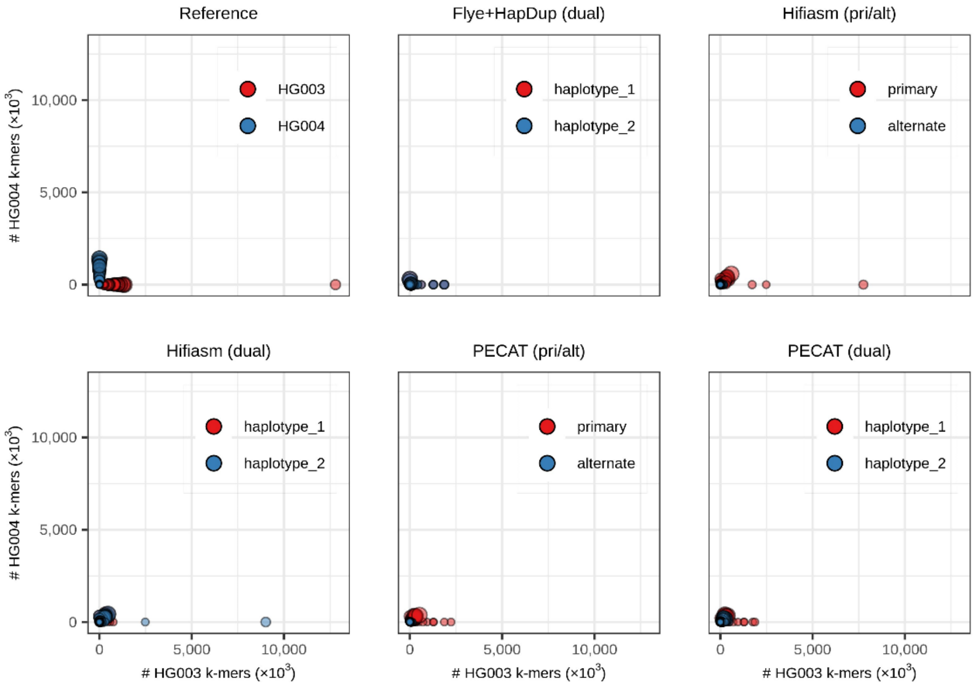


**Supplementary Figure 24. Haplotype-specific k-mer blob plots of the HG002 reference genome and assemblies by different methods from PacBio HiFi** reads. pri/alt or dual represents that the assembly is in the primary/alternate format or the dual assembly format. Each blob corresponds to a contig. The coordinate of the blob gives the count of the parental specific k-mers in the contig, where k is 21. Blob size is proportional to contig length.

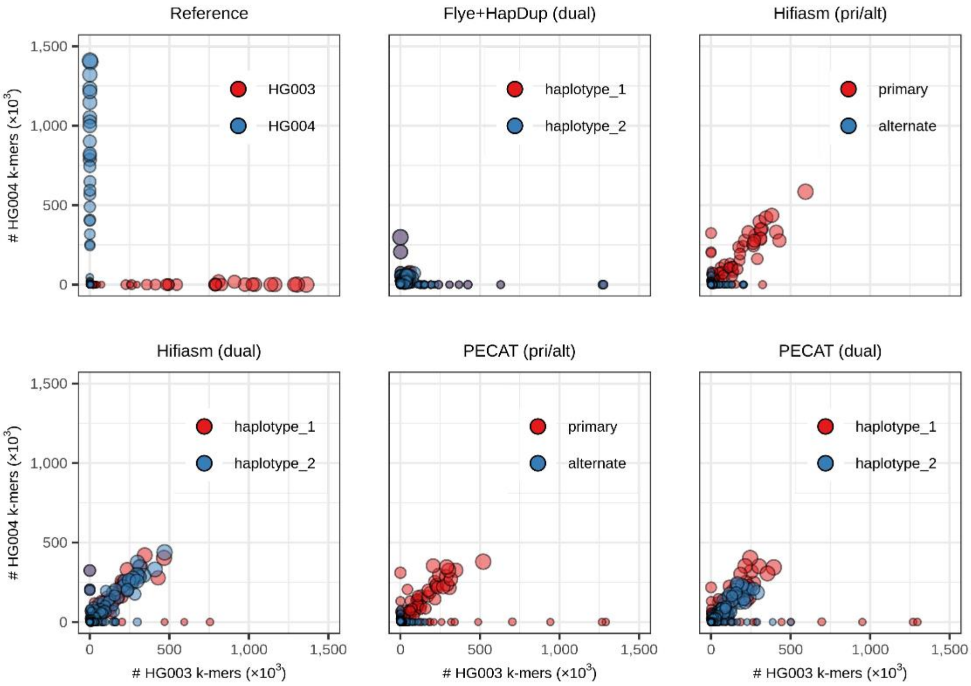


**Supplementary Figure 25**. Enlarged view of **Supplementary Figure 24**.

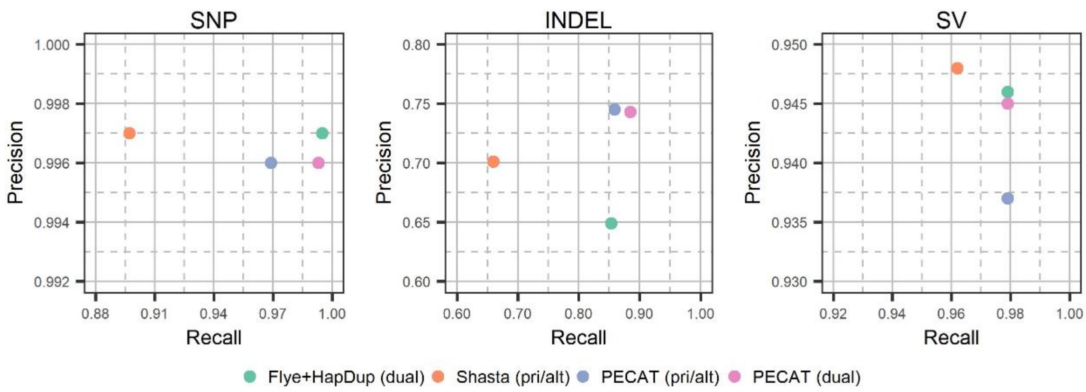


**Supplementary Figure 26**. Precisions and recalls of small variants (SNP, INDEL) and structural variants (SV) in HG002 assemblies from Nanopore R10 duplex reads. 'pri/alt' represents primary/alternate format. 'dual' represents dual assembly format.

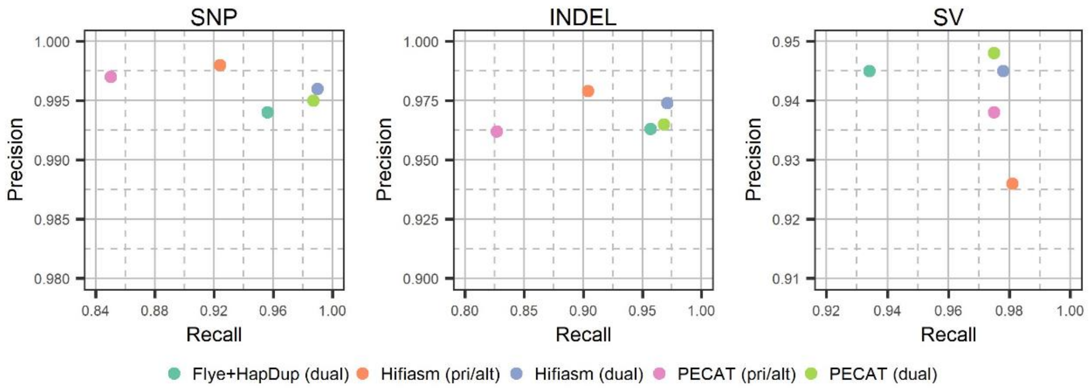


**Supplementary Figure 27**. Precisions and recalls of small variants (SNP, INDEL) and structural variants (SV) in HG002 assemblies from PacBio HiFi reads. 'pri/alt' represents primary/alternate format. 'dual' represents dual assembly format.

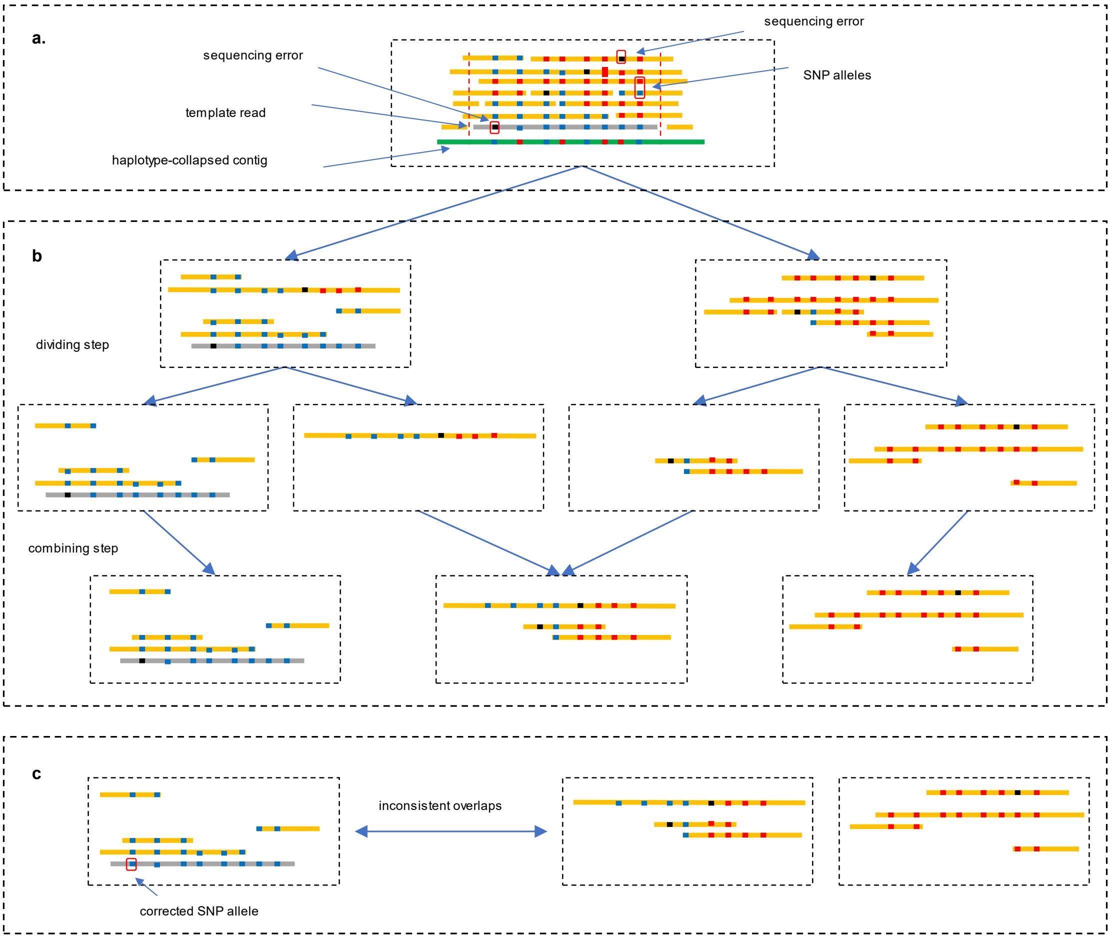


**Supplementary Figure 28. Illustration of identifying inconsistent overlaps.** (**a**) The query reads are collected for each template read. (**b**) A divide-then-combine strategy is used to cluster reads. (**c**) SNP alleles in the template reads are corrected. The inconsistent overlaps between the template read and the query reads are identified.

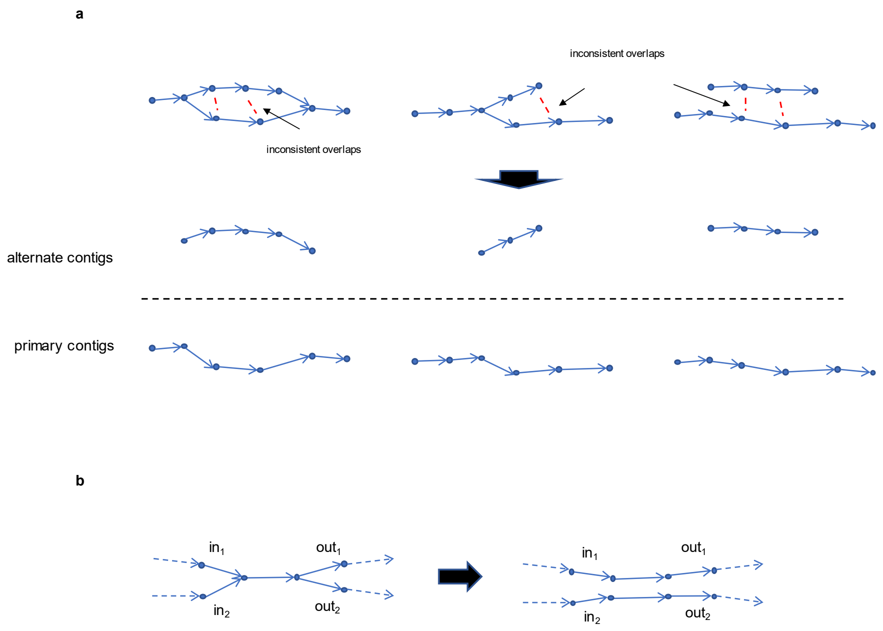


**Supplementary Figure 29. Illustration of generating two sets of contigs. (a)** Three structures are found in the string graph. The two paths are identified in each structure, corresponding to the primary contig and the alternate contig respectively. The red dashed lines mean that there are inconsistent overlaps between the reads in the two paths. **(b)** Phasing two adjacent bubble structures for dual assembly format.
