## Supplementary Notes for "*de novo* diploid genome assembly using long noisy reads"

Jianxin Wang

### Supplementary Note 1: The details of the datasets

The dataset of *S. cerevisiae* (SK×Y12) is a pseudo-diploid dataset combining two haploid yeast strains SK1 and Y12. For each yeast strain, we downsampled 100X data. For *D. melanogaster* (ISO1×A4), we used the same data as Sergey's paper[1] for a fair comparison. The dataset is available at https://obj.umiacs.umd.edu/marbl_publications/hicanu/index.html. *A. thaliana* (Col-0 × C24) was generated using our in-house sequencing.

### *A. thaliana* (Col-0 × C24)

#### Plant materials and growth conditions

The Arabidopsis Columbia-0 (Col-0) and C24 ecotypes were used as the parental lines, and the F1 hybrids between Col-0 and C24 were generated as described previously[2]. Seeds were surface sterilized and then grown on Murashige and Skoog (MS) medium agar plates (½ MS salts, 2% sucrose, 1.2% agar, pH 5.7) for 7 days. The 7d-old seedlings were transplanted on soil pots and grown for three weeks at 22 ± 1℃ with a 16-h-light/8-h-dark photoperiod. The whole seedlings were then harvested for subsequent analysis. The authenticity of F1 progenies from Col and C24 hybridization was genotyped as described previously[2].

#### DNA extraction and purification

Samples were collected, and high molecular weight genomic DNA was prepared by the CTAB method and followed by purification with a QIAGEN® Genomic kit (Cat#13343, QIAGEN) for regular sequencing, according to the standard operating procedure provided by the manufacturer. Ultra-long DNA was extracted by the SDS method without a purification step to sustain the length of DNA. The DNA degradation and contamination of the extracted DNA were monitored on 1% agarose gels. DNA purity was then detected using NanoDrop™ One UV-Vis spectrophotometer (Thermo Fisher Scientific, USA), of which OD260/280 ranges from 1.8 to 2.0 OD 260/230 is between 2.0-2.2. At last, DNA concentration was further measured by Qubit® 4.0 Fluorometer (Invitrogen, USA).

**Nanopore whole genome sequencing and base-calling**

A total amount of 3-4 μ g DNA per sample was used as input material for the ONT library preparations. After the sample was qualified, size-select of long DNA fragments was performed using the PippinHT system (Sage Science, USA). Next, the ends of DNA fragments were repaired, and A-ligation reactions were conducted with NEBNext Ultra II End Repair/dA-tailing Kit (Cat# E7546). The adapter in the SQK-LSK109 (Oxford Nanopore Technologies, UK) was used for further ligation reaction and the DNA library was measured by Qubit® 4.0 Fluorometer (Invitrogen, USA). About 700ng DNA library was constructed and performed on a Nanopore PromethION sequencer instrument (Oxford Nanopore Technologies, UK) at the Genome Center of Grandomics (Wuhan, China). The raw data, collected in this experiment, were obtained as fast5 files after the conversion of electrical signals into base calls via Guppy v5.0.16 (Oxford Nanopore Technologies).

### Simulated data

We first generated paternal and maternal haplotype genomes based on the chromosome chrIV of the yeast SK1 genome. We ran the command to randomly change the bases so that the heterozygosity rate of the dataset was 0.01, 0.005, 0.001, 0.0005, and 0.0001 respectively.

```
Python3 $PECAT/scripts/fxtool.py fx_gen_dipref -r sk1.fa -c chrIV -f
fa_sk1_chrIV.fa -m mo_sk1_chrIV.fa -s variants_sites.txt -p 0.01
```

Next, we ran pbsim2 (eeb5a19)[3] to simulate PacBio CLR reads and Nanopore reads with the following commands.

```
pbsim --depth 30 --prefix fa --hmm_model $PBSIM2/data/P6C4.model fa_sk1_chrIV.fa --length-mean 25000 --id-prefix F
```

```
pbsim --depth 30 --prefix mo --hmm_model $PBSIM2/data/P6C4.model mo_sk1_chrIV.fa --length-mean 25000 --id-prefix M
```

```
pbsim --depth 30 --prefix fa --hmm_model $PBSIM2/data/R103.model fa_sk1_chrIV.fa --length-mean 25000 --id-prefix F
```

```
pbsim --depth 30 --prefix mo --hmm_model $PBSIM2/data/R103.model mo_sk1_chrIV.fa --length-mean 25000 --id-prefix M
```

### Supplementary Note 2: Commands for error correction and assembly

#### Commands for Canu and Purge_dups

Canu (v2.1)[4] was run with the following commands for PacBio CLR reads and Nanopore reads:

```
canu -p $SPECIES -d $SPECIES genomeSize=$GENOME_SIZE corOutCoverage=100 -correct -(pacbio|nanopore) $READS maxThreads=$THREADS
canu -p $SPECIES -d $SPECIES genomeSize=$GENOME_SIZE -corrected -(pacbio|nanopore) ./$SPECIES/$SPECIES.correctedReads.fasta.gz batOptions="-dg 6 -db 6 -dr 1 -ca 500 -cp 50" maxThreads=$THREADS
```

Purge_dups (v1.2.5)[5] was run with the following commands to partition the contigs generated by other assemblers to primary and alternate contigs.

```
minimap2 -x map-pb $CONTIGS $READS | gzip -c - > rd2ctg.paf.gz
pbcstat rd2ctg.paf.gz
calcuts PB.stat > cutoffs 2>calcults.log
split_fa $CONTIGS > contigs.split.fasta
minimap2 -xasm5 -DP contigs.split.fasta contigs.split.fasta | gzip -c - > ctg2ctg.paf.gz
```

```
purge_dups -2 -T cutoffs -c PB.base.cov ctg2ctg.paf.gz > dups.bed 2>
purge_dups.log
get_seqs dups.bed $CONTIGS
```

### Commands for FALCON and FALCON-Unzip

FALCON (1.8.1)[6] was run with the following parameters to assemble reads and FACON-Unzip (1.3.7)[6] was run with default parameters.

Parameters for *S. cerevisiae* (SK1×Y12):

```
genome_size=12000000
length_cutoff = 4000
length_cutoff_pr = 4000
pa_daligner_option = -e0.75 -l1800 -k18 -h240 -w8 -s100
ovlp_daligner_option = -k24 -h60 -e.96 -l1500 -s100
pa_HPCdaligner_option = -v -B128 -M24
ovlp_HPCdaligner_option = -v -B128 -M24
pa_DBsplit_option = -a -x500 -s50
ovlp_DBsplit_option = -s50
falcon_sense_option = --output_multi --min_idt 0.70 --min_cov 4 --
max_n_read 400 --n_core 24
overlap_filtering_setting = --max_diff 120 --max_cov 150 --min_cov 4
--n_core 2
falcon_sense_skip_contained = False
```

Parameters for *A. thaliana* (Col-0×Cvi-0):

```
genome_size=130000000
length_cutoff = 4000
length_cutoff_pr = 4000
pa_daligner_option = -e0.75 -l1800 -k18 -h240 -w8 -s100
ovlp_daligner_option = -k24 -h750 -e.96 -l1500 -s100
pa_HPCdaligner_option = -v -B128 -M24
```

```
ovlp_HPCdaligner_option = -v -B128 -M24
pa_DBsplit_option = -a -x500 -s400
ovlp_DBsplit_option = -s400
falcon_sense_option = --output_multi --min_idt 0.70 --min_cov 4 --max_n_read 400 --n_core 24
overlap_filtering_setting = --max_diff 80 --max_cov 120 --min_cov 4 --n_core 24
falcon_sense_skip_contained = False
```

Parameters for *D. melanogaster* (ISO1×A4):

```
genome_size=140000000
length_cutoff = 10000
length_cutoff_pr = 10000
pa_daligner_option = -e0.75 -l1200 -k18 -h480 -w8 -s100
ovlp_daligner_option = -k24 -h480 -e.95 -l1500 -s100
pa_HPCdaligner_option = -v -B128 -M24
ovlp_HPCdaligner_option = -v -B128 -M24
pa_DBsplit_option = -x500 -s400
ovlp_DBsplit_option = -s400
falcon_sense_option = --output_multi --min_idt 0.70 --min_cov 4 --max_n_read 200 --n_core 24
overlap_filtering_setting = --max_diff 1000 --max_cov 1000 --min_cov 2 --n_core 24
falcon_sense_skip_contained = False
```

### Commands for MECAT2

MECAT2 (f54c542)[7] was run with the command “mecat.pl correct cfg” for correcting PacBio CLR reads. The parameter file is shown below:

```
PROJECT=$SPECIES
RAWREADS= $READS
```

```
GENOME_SIZE= $GENOME_SIZE
THREADS=THREADS
MIN_READ_LENGTH=2000
CNS_OVLP_OPTIONS="-kmer_size 13"
CNS_PCAN_OPTIONS="-p 100000 -k 100"
CNS_OPTIONS=""
CNS_OUTPUT_COVERAGE=80
TRIM_OVLP_OPTIONS="-skip_overhang"
TRIM_PM4_OPTIONS="-p 100000 -k 100"
TRIM_LCR_OPTIONS=""
TRIM_SR_OPTIONS=""
ASM_OVLP_OPTIONS=""
FSA_OL_FILTER_OPTIONS="--max_overhang=-1 --min_identity=-1"
FSA_ASSEMBLE_OPTIONS=""
```

To speed up correcting *B.taurus* (Angus × Brahman) reads, we adjusted the parameters as shown below:

```
CNS_OVLP_OPTIONS="-kmer_size 17"
```

### Commands for NECAT

NECAT (47c6c23)[8] was run with the command "necat.pl correct cfg' for correcting Nanopore reads. The parameter file is shown below:

```
PROJECT= $SPECIES
ONT_READ_LIST=$READS
GENOME_SIZE=$GENOME_SIZE
THREADS=$THREADS
MIN_READ_LENGTH=3000
PREP_OUTPUT_COVERAGE=80
OVLP_FAST_OPTIONS=-n 500 -z 20 -b 2000 -e 0.5 -j 0 -u 1 -a 1000
OVLP_SENSITIVE_OPTIONS=-n 500 -z 10 -e 0.5 -j 0 -u 1 -a 1000
```

```
CNS_FAST_OPTIONS=-a 2000 -x 4 -y 12 -l 1000 -e 0.5 -p 0.8 -u 0
CNS_SENSITIVE_OPTIONS=-a 2000 -x 4 -y 12 -l 1000 -e 0.5 -p 0.8 -u 0
TRIM_OVLP_OPTIONS=-n 100 -z 10 -b 2000 -e 0.5 -j 1 -u 1 -a 400
ASM_OVLP_OPTIONS=-n 100 -z 10 -b 2000 -e 0.5 -j 1 -u 0 -a 400
NUM_ITER=2
CNS_OUTPUT_COVERAGE=80
CLEANUP=1
USE_GRID=false
GRID_NODE=0
GRID_OPTIONS=
SMALL_MEMORY=0
FSA_OL_FILTER_OPTIONS=
FSA_ASSEMBLE_OPTIONS=
FSA_CTG_BRIDGE_OPTIONS=
POLISH_CONTIGS=true
```

### Commands for Flye + HapDup

Flye (2.9)[9] was run to assemble reads: *A. thaliana* (Col-0×C24), *B. taurus* (Bison × Simmental), and HG002. Flye was run with the command:

```
flye --nano-raw|--nano-hq|--pacbio-hifi $READS --out-dir $SPECIES --threads 48 --genome-size $GENOME_SIZE
```

For *B. taurus* (Bison × Simmental) and HG002, we added parameter `--asm-coverage 50`.

Then, HapDup (0.5 or 0.12)[10,11]was run with the command:

```
minimap2 assembly.fasta $READS -a -x map-ont  -t 48 | samtools sort -@4 -m 4G  > rd2ctg.sorted.bam
samtools index -@ 48 rd2ctg.sorted.bam
HD_DIR=`pwd`
```

```
singularity exec --bind $HD_DIR hapdup_0.5.sif hapdup --assembly $HD_DIR/assembly.fasta --bam $HD_DIR/rd2ctg.sorted.bam --out-dir $HD_DIR/hapdup -t 48 --rtype ont
```

where `assembly.fasta` is the assembly generated by Flye. HapDup (0.12) was run assemble Nanopore R10 sequencing (ultra-long), Nanopore R10 duplex sequencing and PacBio HiFi sequencing reads.

### Commands for Shasta

Shasta (0.9.0 or 0.11.1)[12] was run with the following command for Nanopore reads *A. thaliana* (Col-0×C24), *B. taurus* (Bison × Simmental), and HG002.

```
shasta-Linux-0.9.0 --input $READS --config (Nanopore-Phased-Jan2022|Nanopore-UL-Phased-Jan2022) --threads $THREADS
```

Shasta (v0.11.1) was run to assemble Nanopore R10 sequencing (ultra-long) and Nanopore R10 duplex sequencing reads. Then, the following command was used to connect the haplotigs in "Assembly-Phased.fasta" file to generate primary/alternate-style contigs.

```
python3 $PECAT/fxtools.py fx_split_shasta Assembly-Phased.fasta primary.fasta alternate.fasta --min-length 500
```

### Commands for PECAT

PECAT (v0.0.2 or v0.0.3) was run to assemble genomes. PECAT (v0.0.3) was run to assemble Nanopore R10 sequencing (ultra-long), Nanopore R10 duplex sequencing and PacBio HiFi sequencing reads. PECAT was run with the command 'pecat.pl correct cfg' for correcting and the command 'pecat.pl unzip cfg' for assembling.

Parameters for *S. cerevisiae* (SK1×Y12):

```
project=yeast
reads= $READS
genome_size=12000000
threads=48
cleanup=1
grid=local

prep_min_length=3000
```

```
prep_output_coverage=80

corr_iterate_number=1
corr_block_size=4000000000
corr_filter_options=--
filter0=al=2000:alr=0.5:aalr=0.5:oh=1000:ohr=0.1
corr_correct_options=--score=weight:lc=10  --aligner  diff  --filter1
oh=1000
corr_rd2rd_options=-x ava-pb
corr_output_coverage=80

align_block_size=4000000000
align_rd2rd_options=-X -g3000 -w30 -k19 -m100 -r500
align_filter_options=--
filter0=l=5000:al=5000:aal=6000:aalr=0.5:oh=1000:ohr=0.1           --
task=extend --filter1=oh=100:ohr=0.01

asm1_assemble_options=--reducer1 spur:length=1000:nodesize=3

phase_method=0
phase_rd2ctg_options=-x map-pb -c -p 0.5 -r 1000
phase_use_reads=1
phase_phase_options= --phase_options icr=0.2
phase_filter_options=

asm2_assemble_options=--reducer1    spur:length=1000:nodesize=3    --
contig_format dual,prialt

polish_use_reads=1
polish_map_options = -x asm20
polish_cns_options =
```

*A. thaliana* (Col-0×Cvi-0) and *D. melanogaster* (ISO1×A4)

```
project= $SPCIES
reads= $READS
genome_size=$GENOME_SIZE
threads=48
cleanup=1
grid=local

prep_min_length=3000
prep_output_coverage=80

corr_iterate_number=1
corr_block_size=4000000000
```

```
corr_filter_options=--
filter0=:al=5000:alr=0.5:aal=8000:aalr=0.5:oh=2000:ohr=0.2
corr_correct_options=--score=weight:lc=10  --aligner  diff  --filter1
oh=1000:ohr=0.01
corr_rd2rd_options=-x ava-pb
corr_output_coverage=80

align_block_size=4000000000
align_rd2rd_options=-X -g3000 -w30 -k19 -m100 -r500
align_filter_options=--
filter0=l=3000:al=3000:alr=0.5:aalr=0.5:oh=1000:ohr=0.1 --task=extend
--filter1=oh=100:ohr=0.01

asm1_assemble_options=--max_trivial_length 10000

phase_rd2ctg_options=-x map-pb -c -p 0.5 -r 1000
phase_use_reads=1
phase_phase_options= --phase_options icr=0.2
phase_filter_options= --threshold=1000

asm2_assemble_options=--max_trivial_length    10000    --contig_format
dual,prialt

polish_use_reads=1
polish_map_options = -x asm20
polish_cns_options =
```

Parameters for *B. taurus* (Angus×Brahman):

```
project=cattle
reads= $READS
genome_size= 2700000000
threads=48
grid=local
cleanup=1

prep_min_length=3000
prep_output_coverage=80

corr_iterate_number=1
corr_block_size=8000000000
corr_filter_options=--
filter0=:al=5000:alr=0.5:aal=8000:aalr=0.5:oh=2000:ohr=0.2
corr_correct_options=--score=weight:lc=10 --aligner diff --filter1
oh=1000:ohr=0.01 --candidate n=300:f=20
corr_rd2rd_options=-x ava-pb -f 0.005 -I 20G
```

```
corr_output_coverage=80

align_block_size=4000000000
align_rd2rd_options=-X -g3000 -w30 -k19 -m100 -r500 -I 20G -f 0.005
align_filter_options=--
filter0=l=3000:al=3000:alr=0.5:aalr=0.5:oh=1000:ohr=0.1 --task=extend
--filter1=oh=100:ohr=0.01

asm1_assemble_options= --max_trivial_length 10000

phase_rd2ctg_options=-x map-pb -c -p 0.5 -r 1000
phase_use_reads=1
phase_phase_options= --phase_options icr=0.2
phase_filter_options= --threshold=1000

asm2_assemble_options= --max_trivial_length 10000 --contig_format
dual,prialt

polish_use_reads=1
polish_map_options = -x asm20
polish_filter_options = --filter0 oh=1000:ohr=0.1
polish_cns_options =
```

Parameters for *A. thaliana* (Col-0 × C24):

```
project= arab
reads= $READS
genome_size= 130000000
threads=48
cleanup=1
grid=local

prep_min_length=3000
prep_output_coverage=80

corr_iterate_number=1
corr_block_size=4000000000
corr_filter_options=--
filter0=l=5000:al=2500:alr=0.5:aal=5000:oh=3000:ohr=0.3
corr_correct_options=--score=weight:lc=10 --aligner edlib --filter1
oh=1000:ohr=0.01
corr_rd2rd_options=-x ava-ont -k19
corr_output_coverage=80

align_block_size=4000000000
```

```
align_rd2rd_options=-X -g3000 -w30 -k19 -m100 -r500 -f 0.001
align_filter_options=--
filter0=l=5000:aal=6000:aalr=0.5:oh=3000:ohr=0.3 --task=extend --
filter1=oh=300:ohr=0.03
asm1_assemble_options=--max_trivial_length 10000

phase_method=2
phase_rd2ctg_options=-x map-ont -c -p 0.5 -r 1000
phase_phase_options= --coverage lc=30 --phase_options
icr=0.1:icc=8:sc=10
phase_use_reads=1
phase_filter_options= --threshold=1000

phase_clair3_command = singularity exec -B `pwd -P`:`pwd -P`
clair3_v0.1-r12.sif /opt/bin/run_clair3.sh
phase_clair3_options=--platform=ont --
model_path=/opt/models/ont_guppy5/  --include_all_ctgs
phase_clair3_rd2ctg_options=-x map-ont -c -p 0.5 -r 1000
phase_clair3_phase_options= --coverage lc=30 --phase_options
icr=0.1:icc=6:sc=10 --filter i=70
phase_clair3_use_reads=0
phase_clair3_filter_options= --threshold=2500 --rate 0.05

asm2_assemble_options=--max_trivial_length 10000 --contig_format
dual,prialt

polish_map_options = -x map-ont -k19 -w10 -I 10g
polish_use_reads=0
polish_cns_options =

polish_medaka = 1
polish_medaka_command =singularity exec -B `pwd -P`:`pwd -P`
medaka_v1.7.2.sif medaka
polish_medaka_map_options = -x map-ont  -k19 -w10 -I 10g
polish_medaka_cns_options = --model r941_prom_sup_g507
```

Parameters for *B. taurus* (Bison × Simmental):

```
project= cattle
reads= $READS
genome_size= 2700000000
threads=48
cleanup=1
grid=local
prep_min_length=3000
prep_output_coverage=80
```

```
corr_iterate_number=1
corr_block_size=8000000000
corr_filter_options=--
filter0=l=5000:al=2500:alr=0.5:aal=8000:oh=3000:ohr=0.3
corr_correct_options=--score=weight:lc=16 --aligner edlib --filter1
oh=1000:ohr=0.01 --candidate n=400:f=20
corr_rd2rd_options=-x ava-ont -f 0.005 -I 10G
corr_output_coverage=80
align_block_size=12000000000
align_rd2rd_options=-X -g3000 -w30 -k19 -m100 -r500 -I 10G -f 0.005
align_filter_options=--
filter0=l=5000:aal=6000:aalr=0.5:oh=3000:ohr=0.3 --task=extend --
filter1=oh=300:ohr=0.03
asm1_assemble_options=--max_trivial_length 1000000

phase_method=2
phase_rd2ctg_options=-x map-ont -c -p 0.5 -r 1000
phase_use_reads=1
phase_phase_options= --coverage lc=30 --phase_options
icr=0.1:icc=8:sc=10
phase_filter_options = --threshold 1000

phase_clair3_command = singularity exec --containall -B `pwd -P`:`pwd
-P` clair3_v0.1-r12.sif /opt/bin/run_clair3.sh
phase_clair3_options=--platform=ont --
model_path=/opt/models/ont_guppy5/  --include_all_ctgs
phase_clair3_rd2ctg_options=-x map-ont -c -p 0.5 -r 1000
phase_clair3_use_reads=0
phase_clair3_phase_options= --coverage lc=30 --phase_options
icr=0.1:icc=6:sc=10 --filter i=70
phase_clair3_filter_options = --threshold 2500 --rate 0.05

asm2_assemble_options=--max_trivial_length 1000000 --contig_format
dual,prialt
polish_map_options = -x map-ont -w10 -k19
polish_use_reads=0
polish_filter_options=--filter0 oh=2000:ohr=0.2:aalr=0.5
polish_cns_options =

polish_medaka = 1
polish_medaka_command = singularity exec -B `pwd -P`:`pwd -P`
medaka_v1.7.2.sif medaka
polish_medaka_map_options = -x map-ont -w10 -k19
polish_medaka_cns_options = --model r941_prom_sup_g507
```

Parameters for HG002 (ONT R9 UL):

```
project= human
reads= $READS
genome_size= 3000000000
threads=48
cleanup=1
grid=local
prep_min_length=3000
prep_output_coverage=80
corr_iterate_number=1
corr_block_size=4000000000
corr_filter_options=--
filter0=l=5000:al=2500:alr=0.5:aal=8000:oh=3000:ohr=0.3
corr_correct_options=--score=weight:lc=10 --aligner edlib --filter1
oh=1000:ohr=0.01 --candidate n=600:f=30
corr_rd2rd_options=-x ava-ont -f 0.005 -I 10G
corr_output_coverage=80

align_block_size=12000000000
align_rd2rd_options=-X -g3000 -w30 -k19 -m100 -r500 -I 10G -f 0.002
align_filter_options=--
filter0=l=5000:aal=6000:aalr=0.5:oh=3000:ohr=0.3 --task=extend --
filter1=oh=300:ohr=0.03
asm1_assemble_options=--max_trivial_length 10000

phase_method=2
phase_rd2ctg_options=-x map-ont -c -p 0.5 -r 1000
phase_use_reads=1
phase_phase_options= --coverage lc=30 --phase_options
icr=0.1:icc=8:sc=10
phase_filter_options = --threshold 1000

phase_clair3_use_reads=0
phase_clair3_command = singularity exec -B `pwd -P`:`pwd -P`
clair3_v0.1-r12.sif /opt/bin/run_clair3.sh
phase_clair3_options=--platform=ont --
model_path=/opt/models/ont_guppy5/  --include_all_ctgs
phase_clair3_rd2ctg_options=-x map-ont -c -p 0.5 -r 1000
phase_clair3_phase_options= --coverage lc=30 --phase_options
icr=0.1:icc=3:sc=10 --filter i=70
phase_clair3_filter_options = --threshold 2500 --rate 0.05

asm2_assemble_options=--reducer0 "best:cmp=2,0.1,0.1|phase:sc=3" --
contig_format dual,prialt --min_identity 0.98
```

```
polish_map_options = -x map-ont -w10 -k19 -I 10g
polish_use_reads=0
polish_filter_options=--filter0 oh=2000:ohr=0.2:aalr=0.5
polish_cns_options =

polish_medaka = 1
polish_medaka_command = singularity exec -B `pwd -P`:`pwd -P`
medaka_v1.7.2.sif medaka
polish_medaka_map_options = -x map-ont -w10 -k19
polish_medaka_cns_options = --model r941_prom_sup_g507
```

Parameters for HG002 (ONT R10 UL):

```
project= human
reads= $READS
genome_size= $GENOME_SIZE
threads=48
cleanup=0
compress=0
grid=auto
prep_min_length=3000
prep_output_coverage=80
corr_iterate_number=1
corr_block_size=4000000000
corr_filter_options=--
filter0=l=5000:al=2500:alr=0.5:aal=8000:oh=3000:ohr=0.3
corr_correct_options=--score=weight:lc=10 --aligner
edlib:bs=1000:mc=6  --min_coverage 4 --filter1 oh=1000:ohr=0.01 --
candidate n=600:f=30 --min_identity 90 --min_local_identity 80
corr_rd2rd_options=-x ava-ont -f 0.005 -I 10G
corr_output_coverage=60
align_block_size=12000000000
align_rd2rd_options=-X -g3000 -w30 -k19 -m100 -r500 -I 10G -f 0.002
align_filter_options=--
filter0=l=5000:aal=6000:aalr=0.5:oh=3000:ohr=0.3 --task=extend --
filter1=oh=300:ohr=0.03 --min_identity 0.95
asm1_assemble_options=--max_trivial_length 10000

phase_method=2
phase_rd2ctg_options=-x map-ont -w 10 -k19  -c -p 0.5 -r 1000 -I 10G
-K 8G
phase_use_reads=1
phase_phase_options= --coverage lc=20 --phase_options
icr=0.1:icc=8:sc=10
phase_filter_options = --threshold 1000
```

```
phase_clair3_command = singularity exec --containall -B `pwd -P`:`pwd
-P` -B /tmp:/tmp clair3_v0.1-r12.sif /opt/bin/run_clair3.sh
phase_clair3_rd2ctg_options=-x map-ont -w10 -k19 -c -p 0.5 -r 1000 -I
10G -K 8G
phase_clair3_use_reads=0
phase_clair3_phase_options= --coverage lc=20 --phase_options
icr=0.1:icc=3:sc=10 --filter i=90
phase_clair3_filter_options = --threshold 2500 --rate 0.05
phase_clair3_options=--platform=ont --
model_path=/opt/models/r941_prom_sup_g5014  --include_all_ctgs

asm2_assemble_options=--reducer0 "best:cmp=2,0.1,0.1|phase:sc=3" --
contig_format prialt,dual

polish_map_options = -x map-ont -w10 -k19 -I 10g -K 8G -a
polish_cns_options =
polish_use_reads=0
polish_filter_options=--filter0 oh=2000:ohr=0.2:i=96

polish_medaka = 1
polish_medaka_command = singularity exec --containall -B `pwd -
P`:`pwd -P` medaka_v1.7.2.sif medaka
polish_medaka_map_options = -x map-ont -w10 -k19 -I 10g -K 8G
polish_medaka_cns_options = --model r1041_e82_260bps_sup_g632
polish_medaka_filter_options=--filter0 oh=2000:ohr=0.2:i=96
```

Parameters for HG002 (ONT R10 Duplex):

```
project= human
reads= $READS
genome_size= $GNOME_SIZE
threads=40
cleanup=0
grid=auto
prep_min_length=3000
prep_output_coverage=60
corr_iterate_number=1
corr_block_size=4000000000
corr_correct_options=--score=weight:lc=8 --aligner diff:s=500 --
min_coverage 1 --filter1 oh=1000:ohr=0.01 --min_identity 95 --
min_local_identity 90 --candidate n=600:f=30
corr_filter_options=--
filter0=l=5000:al=2500:alr=0.5:aal=5000:oh=3000:ohr=0.3
corr_rd2rd_options=-X -g3000 -w30 -k19 -m100 -r500 -f 0.002 -K8G -I
8G
```

```
corr_output_coverage=60
align_block_size=4000000000
align_rd2rd_options=-X -g3000 -w30 -k19 -m100 -r500 -f 0.002 -K8G -I 8G
align_filter_options=--filter0=l=5000:aal=6000:aalr=0.5:oh=3000:ohr=0.3 --task=extend --filter1=oh=50:ohr=0.01  --aligner diff:s=100 --min_identity 0.90
asm1_assemble_options= --min_identity 0.99 --min_coverage 1

phase_method=2
phase_rd2ctg_options=-x map-ont -w10 -k19 -c -p 0.5 -r 1000 -I 10G
phase_use_reads=1
phase_phase_options= --coverage lc=8 --phase_options icr=0.02:icc=3:sc=4 --filter=i=95.00:alr=0.80:oh=100:ohr=0.01:ilid=100

phase_clair3_command=singularity exec --containall -B `pwd -P`:`pwd -P` -B /tmp:/tmp clair3_v0.1-r12.sif /opt/bin/run_clair3.sh
phase_clair3_use_reads=0
phase_clair3_options=--platform=ont --model_path=/opt/models/ont_guppy5/  --include_all_ctgs
phase_clair3_rd2ctg_options=-x map-ont -w10 -k19 -c -p 0.5 -r 1000 -I 10G -K 8G
phase_clair3_phase_options=--coverage lc=8 --phase_options icr=0.02:icc=2:sc=4 --filter i=95
phase_clair3_filter_options=--threshold=2500 --rate 0.05

asm2_assemble_options= --reducer0 "best:cmp=2,0.1,0.1|phase:sc=2" --min_identity 0.99 --max_trivial_length 10000 --contig_format dual,prialt --min_coverage 1

polish_map_options=-x map-ont -w10 -k19 -I 10G -K 8G -a
polish_filter_options=--filter0 oh=1000:ohr=0.1:i=98
polish_cns_options=
polish_medaka=1
polish_medaka_command= singularity exec --containall -B `pwd -P`:`pwd -P` medaka_v1.7.2.sif medaka
polish_medaka_map_options=-x map-ont -w10 -k19 -I 10G -K 8G
polish_medaka_cns_options = --model r1041_e82_400bps_sup_g615
polish_medaka_filter_options=--filter0 oh=1000:ohr=0.1:i=98
```

Parameters for HG002 (PacBio HiFi):

```
project= human
reads= $READS
genome_size=$GENOME_SIZE
```

```
threads=48
cleanup=0
grid=auto
prep_min_length=3000
prep_output_coverage=60
corr_iterate_number=1
corr_block_size=4000000000
corr_correct_options=--score=weight:lc=8 --aligner diff:s=100 --
min_coverage 1 --filter1 oh=100 --min_identity 96 --
min_local_identity 95
corr_filter_options=--
filter0=l=5000:al=2500:alr=0.5:aal=5000:oh=3000:ohr=0.3
corr_rd2rd_options=-X -g3000 -w30 -k19 -m100 -r500 -f 0.002 -K8G -I
8G
corr_output_coverage=60
align_block_size=4000000000
align_rd2rd_options=-X -g3000 -w30 -k19 -m100 -r500 -f 0.002 -K8G -I
8G
align_filter_options=--
filter0=l=5000:aal=6000:aalr=0.5:oh=3000:ohr=0.3 --task=extend --
filter1=oh=50:ohr=0.01  --aligner diff:s=100 --min_identity 0.90
asm1_assemble_options= --min_identity 0.99 --min_coverage 1

phase_method=2
phase_rd2ctg_options=-x map-hifi  -c -p 0.5 -r 1000 -K8G
phase_use_reads=1
phase_phase_options= --coverage lc=8 --phase_options
icr=0.02:icc=3:sc=4 --
filter=i=95.00:alr=0.80:oh=100:ohr=0.01:ilid=100

phase_clair3_command= singularity exec --containall -B `pwd -P`:`pwd
-P` -B /tmp:/tmp clair3_v0.1-r12.sif /opt/bin/run_clair3.sh
phase_clair3_use_reads=0
phase_clair3_options=--platform=hifi --model_path=/opt/models/hifi  -
-include_all_ctgs
phase_clair3_rd2ctg_options=-x map-hifi  -c -p 0.5 -r 1000 -K8G
phase_clair3_phase_options=--coverage lc=8 --phase_options
icr=0.02:icc=2:sc=4 --filter i=95
phase_clair3_filter_options=

asm2_assemble_options= --reducer0 "best:cmp=2,0.1,0.1|phase:sc=2" --
min_identity 0.99 --max_trivial_length 10000 --contig_format
dual,prialt --min_coverage 1
```

```
polish_map_options=-x map-hifi -I8G -K8G -a
polish_filter_options=--filter0 oh=500:ohr=0.05:i=98
polish_cns_options=
```

In the above parameters, `$READS` is the path of the reads file. `$GENOME_ SIZE` is the genome size of the species.

### Supplementary Note 3: Evaluating heterozygosity.

We used genomescope2.0[13] and jellyfish (2.3.0)[14] to evaluate the heterozygosity of these species with the following commands.

```
    jellyfish count -C -m 21 -t 24 -s 4000000000 *.fastq -o reads.jf
    jellyfish histo -t 24 reads.jf > reads.histo
    genomescope.R -i reads.histo -o out -k 21
```

We used 60X offspring's Illumina short sequences or combined 30X paternal and maternal Illumina short sequences as input data.

### Supplementary Note 4: Evaluating assemblies with merqury

We followed the instructions on the website https://github.com/marbl/merqury and used the following command to build meryl databases.

```
sh $MERQURY/best_k.sh $GENOME_SIZE
meryl k=$k count output f1.meryl f1.fastq.gz
meryl k=$k count output maternal.meryl maternal.fastq.gz
meryl k=$k count output paternal.meryl paternal.fastq.gz
sh $MERQURY/trio/hapmers.sh maternal.meryl paternal.meryl f1.meryl
```

Then, we used the following command to evaluate the assemblies.

```
$MERQURY/merqury.sh    f1.meryl    maternal.meryl    paternal.meryl
(pri|hap1).fasta (alt|hap2).fasta $PREFIX
```

Merqury counted the number of parental-specific k-mers. We calculated the hamming error rate of the assemblies from the output of merqury using the following command.

```
$PECAT/build/bin/fxtools.py fx_hamming_error $PREFIX.hapmers.count
```

### Supplementary Note 5: Evaluating gene completeness

BUSCO (5.2.2)[15] was run to evaluate the gene completeness of assemblies for all species. We used the following script:

```
busco -i $CONTIG -m geno --cpu $THREADS -l $LIBS --offline -o contigs.busco --out_path $XXX
```

where `$CONTIG` was set to one of the assemblies and `$LIBS` was set to the corresponding OrthoDB v10 dataset. We used datasets saccharomycetes, brassicales, diptera, cetartiodactyla, and primates for *S. cerevisiae*, *A. thaliana*, *D. melanogaster*, *B. taurus*, and HG002, respectively. Those datasets can be downloaded from https://busco-data.ezlab.org/v5/data/lineages/.

### Supplementary Note 6: Evaluating assemblies with Pomoxis.

The program assess_assembly in Pomoxis (https://github.com/nanoporetech/pomoxis) was used to evaluate the quality of the assemblies with the following commands:

```
cat paternal_reference.fna maternal_reference.fna > all_reference.fna
assess_assembly -r all_reference.fna -i primary.fasta
assess_assembly -r all_reference.fna -i alternate.fasta
```

### Supplementary Note 7: Evaluating assembly quality with QUAST.

QUAST (5.0.2)[16] run with the following commands:

```
cat paternal_ref.fna maternal_ref.fna > all_ref.fna
cat primary.fasta alternate.fasta > all_ctgs.fasta
quast.py -r all_ref.fna all_ctgs.fasta --min-contig 5000 --large --min-identity 90
```

For *B. taurus* (Angus×Brahman), *B. taurus* (Bison × Simmental), and HG002, the parameter min-contig was set to 50000.

### Supplementary Note 8: Evaluating SNP, INDEL, and SV in HG002 assemblies

We used dipcall (v0.3)[17] and hap.py(v0.3.15)[18] to calculate the precision, recall and F1-score of the small variants (SNP and INDEL) in HG002 assemblies with following commands:

```
run-dipcall
GCA_000001405.15_GRCh38_no_alt_plus_hs38d1_analysis_set.fna
haplotype_1.fasta haplotype_2.fasta -t 48 > prefix.mak
```

```
make -j2 -f prefix.mak
hap.py HG002_GRCh38_GIAB_highconf_CG-Illfb-IllsentieonHC-Ion-10XsentieonHC-SOLIDgatkHC_CHROM1-22_v.3.3.2_highconf_triophased.vcf.gz prefix.dip.vcf.gz -f HG002_GRCh38_GIAB_highconf_CG-Illfb-IllsentieonHC-Ion-10XsentieonHC-SOLIDgatkHC_CHROM1-22_v.3.3.2_highconf_noinconsistent.bed -r GCA_000001405.15_GRCh38_no_alt_plus_hs38d1_analysis_set.fna -o output -engine=vcfeval
```

We used hapdiff (https://github.com/KolmogorovLab/hapdiff) and truvari[19] to calucate the precision, recall, and F1-score of the structural variant (SV) in HG002 assemblies with the following commands:

```
hapdiff.py --reference hs37d5.fa --pat primary.fasta --mat alternate.fasta --out-dir out -t 48
truvari bench -b HG002_SVs_Tier1_v0.6.vcf.gz -c out/hapdiff_phased.vcf.gz -o result_phased --includebed HG002_SVs_Tier1_v0.6.bed -f hs37d5.fa.gz -r 2000 --chunksize 2000 –passonly
```

The standard SV set is available from GIAB[20]. It can be downloaded from at https://ftp-trace.ncbi.nlm.nih.gov/giab/ftp/data/AshkenazimTrio/analysis/NIST_SVs_Integration_v0.6

### References


1. Nurk, S. *et al.* HiCanu: accurate assembly of segmental duplications, satellites, and allelic variants from high-fidelity long reads. *Genome Res.* **30**, 1291–1305 (2020).

2. Zhang, Q. *et al.* The chromatin remodeler DDM1 promotes hybrid vigor by regulating salicylic acid metabolism. *Cell Discov.* **2**, 16027 (2016).

3. Ono, Y., Asai, K. & Hamada, M. PBSIM2: a simulator for long-read sequencers with a novel generative model of quality scores. *Bioinformatics* **37**, 589–595 (2021).

4. Koren, S. *et al.* Canu: scalable and accurate long-read assembly via adaptive *k* -mer weighting and repeat separation. *Genome Res.* **27**, 722–736 (2017).

5. Guan, D. *et al.* Identifying and removing haplotypic duplication in primary genome assemblies. *Bioinformatics* **36**, 2896–2898 (2020).
6. Chin, C.-S. *et al.* Phased diploid genome assembly with single-molecule real-time sequencing. *Nat. Methods* **13**, 1050–1054 (2016).
7. Xiao, C.-L. *et al.* MECAT: fast mapping, error correction, and de novo assembly for single-molecule sequencing reads. *Nat. Methods* **14**, 1072–1074 (2017).
8. Chen, Y. *et al.* Efficient assembly of nanopore reads via highly accurate and intact error correction. *Nat. Commun.* **12**, 60 (2021).
9. Kolmogorov, M., Yuan, J., Lin, Y. & Pevzner, P. A. Assembly of long, error-prone reads using repeat graphs. *Nat. Biotechnol.* **37**, 540–546 (2019).
10. Shafin, K. *et al.* Haplotype-aware variant calling with PEPPER-Margin-DeepVariant enables high accuracy in nanopore long-reads. *Nat. Methods* **18**, 1322–1332 (2021).
11. Kolmogorov, M. *et al. Scalable Nanopore sequencing of human genomes provides a comprehensive view of haplotype-resolved variation and methylation*. http://biorxiv.org/lookup/doi/10.1101/2023.01.12.523790 (2023) doi:10.1101/2023.01.12.523790.
12. Shafin, K. *et al.* Nanopore sequencing and the Shasta toolkit enable efficient de novo assembly of eleven human genomes. *Nat. Biotechnol.* **38**, 1044–1053 (2020).
13. Ranallo-Benavidez, T. R., Jaron, K. S. & Schatz, M. C. GenomeScope 2.0 and Smudgeplot for reference-free profiling of polyploid genomes. *Nat. Commun.* **11**, 1432 (2020).
14. Marçais, G. & Kingsford, C. A fast, lock-free approach for efficient parallel counting of occurrences of k-mers. *Bioinformatics* **27**, 764–770 (2011).

15. Manni, M., Berkeley, M. R., Seppey, M., Simão, F. A. & Zdobnov, E. M. BUSCO Update: Novel and Streamlined Workflows along with Broader and Deeper Phylogenetic Coverage for Scoring of Eukaryotic, Prokaryotic, and Viral Genomes. *Mol. Biol. Evol.* **38**, 4647–4654 (2021).
16. Gurevich, A., Saveliev, V., Vyahhi, N. & Tesler, G. QUAST: quality assessment tool for genome assemblies. *Bioinformatics* **29**, 1072–1075 (2013).
17. Li, H. *et al.* A synthetic-diploid benchmark for accurate variant-calling evaluation. *Nat. Methods* **15**, 595–597 (2018).
18. the Global Alliance for Genomics and Health Benchmarking Team *et al.* Best practices for benchmarking germline small-variant calls in human genomes. *Nat. Biotechnol.* **37**, 555–560 (2019).
19. English, A. C., Menon, V. K., Gibbs, R. A., Metcalf, G. A. & Sedlazeck, F. J. Truvari: refined structural variant comparison preserves allelic diversity. *Genome Biol.* **23**, 271 (2022).
20. Zook, J. M. *et al.* A robust benchmark for detection of germline large deletions and insertions. *Nat. Biotechnol.* (2020) doi:10.1038/s41587-020-0538-8.
